# Brain-wide analysis of the supraspinal connectome reveals anatomical correlates to functional recovery after spinal injury

**DOI:** 10.1101/2021.06.10.447885

**Authors:** Zimei Wang, Adam Romanski, Vatsal Mehra, Yunfang Wang, Benjamin C. Campbell, Gregory A. Petsko, Pantelis Tsoulfas, Murray Blackmore

## Abstract

The supraspinal connectome is essential for normal behavior and homeostasis and consists of numerous sensory, motor, and autonomic projections from brain to spinal cord. Study of supraspinal control and its restoration after damage has focused mostly on a handful of major populations that carry motor commands, with only limited consideration of dozens more that provide autonomic or crucial motor modulation. We now provide an experimental platform and associated web-based resource to rapidly profile the entire supraspinal mesoconnectome in adult mice. Optimized viral labeling, 3D imaging, and registration to a mouse digital neuroanatomical atlas assigned tens of thousands of supraspinal neurons to more than 60 identified regions. We demonstrate the approach’s ability to clarify essential points of topographic mapping between spinal levels, to measure population-specific sensitivity to spinal injury, and to resolve previously unexplained variability in functional recovery. This work will spur progress by broadening understanding and enabling analyses of essential but understudied supraspinal populations.

## Introduction

The brain’s control of the body below the head is achieved largely by axonal inputs to spinal circuits, which then relay commands to the periphery through motor and autonomic output neurons. This supraspinal connectome is highly conserved in mammals, with multiple cell types distributed through the brainstem, midbrain, and motor cortex, each projecting axons to a subset of spinal levels and to selected cell types ^1, 2^. A comprehensive and accessible approach for understanding supraspinal input is crucial to interpret motor and autonomic behavior, and to treat conditions that disrupt descending signals such as stroke, disease, or injury to the spinal cord.

Extensive work spanning almost a century has employed orthograde degeneration, electrical stimulation, and axonal transport tracing methods to characterize the location and function of specific supraspinal neurons in various animals, providing a base of knowledge to understand supraspinal control ^1–5^. Several efforts in rodents have provided more global information by performing retrograde tracing from selected spinal levels, followed by tissue sectioning and manual assignment of labeled cell bodies to regions within the brain ^6–8^. Significant challenges, however, impede the distribution of this foundational knowledge and its application to the study of disease and injury-based disruptions. First, information about the location and types of supraspinal neurons is fragmented and not standardized across numerous studies ^9^. Second, a high level of expertise is required to precisely identify brain regions from two-dimensional tissue series and to build a three-dimensional view of the connections ^10–12^. Third, tissue sectioning and imaging are laborious and time consuming, making experiments to track dynamic changes after injury or disease impractical. Consequently, attention has remained focused on a relatively narrow set of supraspinal populations. For example, in the field of spinal cord injury, the vast majority of studies concern only a handful of descending populations, notably the corticospinal, rubrospinal, raphespinal, and broadly defined reticulospinal ^9, 13–16^. This attention is justified, as these regions serve important motor functions and comprise a majority of descending input ^17^. On the other hand, dozens of additional brain regions also project to the spinal cord, many of which carry essential motor and autonomic commands ^8^. Without tools to easily monitor the totality of the supraspinal connectome, researchers lack even basic information regarding their sensitivity to injury, innate plasticity, or potentially disparate responses to potential pro-regenerative therapies.

Here we present a comprehensive and accessible approach to obtain detailed information about the number and location of descending projection neurons throughout the mouse brain. By combining retrograde viral labeling ^18^, 3D imaging of optically cleared brains ^19^, and registration to standard neuro-anatomical space ^20, 21^, we rapidly identify the specific location of tens of thousands of supraspinal neurons. We present a web-based resource that compares the locations and quantity of supraspinal neurons that project to cervical versus lumbar levels. We further extend this approach to questions related to spinal cord injury by quantifying the region-specific sparing of distinct supraspinal populations in mice that received injuries of graded severity. Interestingly, this analysis revealed correlation between residual brain-spinal cord connectivity and locomotor function in a subset of neural populations. This approach provides an effective tool to disseminate detailed understanding of the supraspinal connectome, and a needed platform to rapidly achieve brain-wide profiling of disruption and restoration of brain-spinal cord connectivity after injury.

## Results

### Optimization of retrograde cell detection in cleared brain tissue

Although tissue clearing techniques offer unprecedented visualization of intact neural structures, overall degradation of fluorescent signal during the process, compounded by optical interference from nearby axon tracts, can significantly limit detection of cell bodies ^19, 22–24^. This detection problem is evident in prior experiments from our lab and others that used spinal injection of AAV2-retro to retrogradely express fluorescent proteins (FPs) in supraspinal neurons ^19, 22^. In tissue sections that spanned the brain we observed highly efficient transduction, as assessed by co-label of viral tdTomato and cholera toxin B mixed as a “gold standard” retrograde marker. tdTomato intensity was variable between different populations, however, and when brains were subjected to 3D tissue clearing it was evident that in dimmer regions only a small fraction of the supraspinal neurons visible in tissue sections were detected in 3D images. This was especially true in the brainstem, an observation echoed by a recent study ^22^. In an effort to improve detection we first replaced tdTomato with mScarlet (mSc), a bright red monomeric fluorescent protein (FP) ^25^. Adult mice received injection of AAV2-retro-mSc to lumbar spinal cord, followed two or four weeks later by 3DISCO-mediated clearing of brains and imaging by light-sheet fluorescence microscopy (LSFM) ^19^. Similar to prior results with tdTomato, however, although CST neurons were labeled brightly (**Figure 1 A,C**) pontine and medullary populations were dimmer, sparse, and partially obscured by the bright pyramidal tracts (**Figure 1 E,F**).

**Figure 1:**
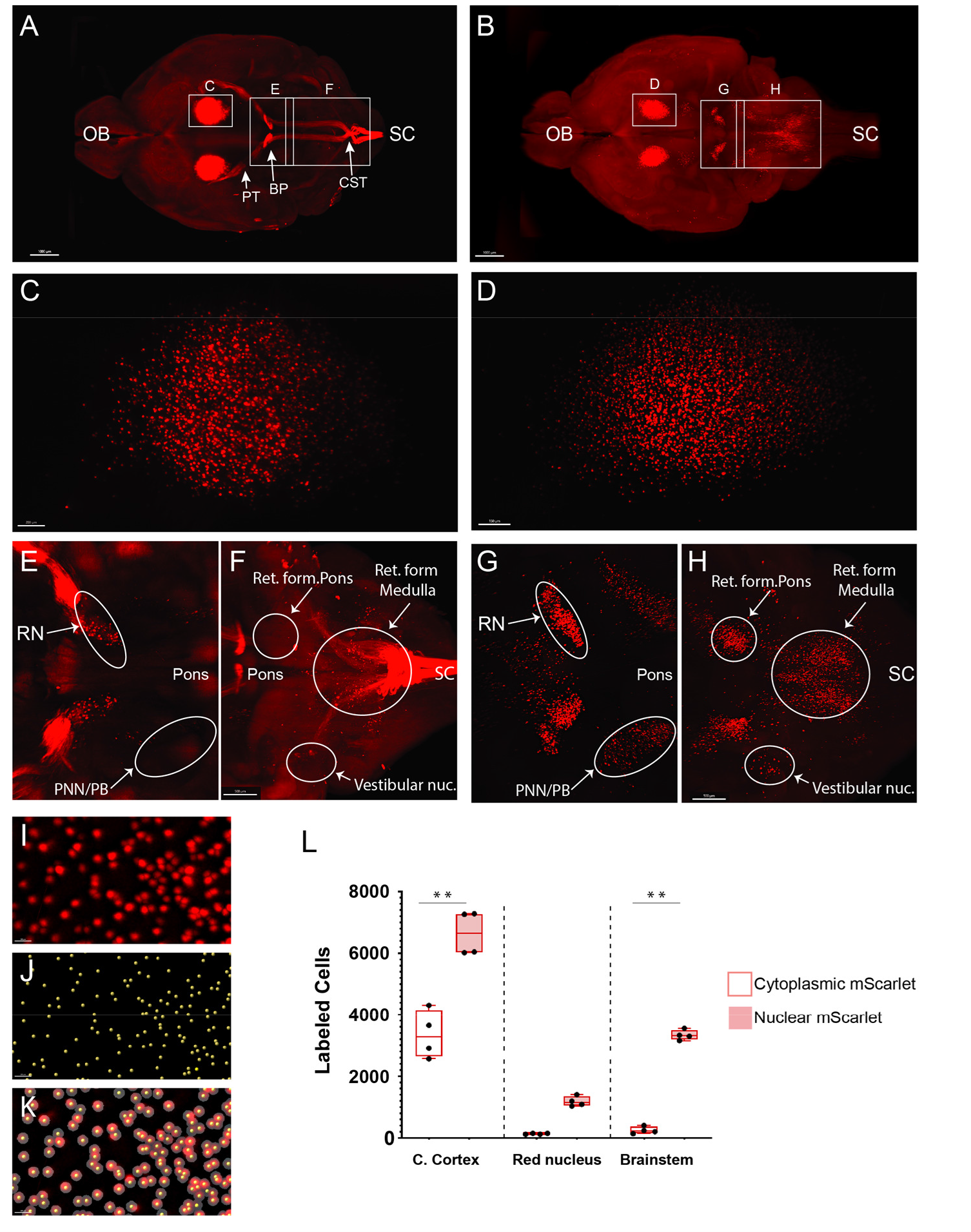
Nuclear localization of retrograde fluorophores enhances detection of supraspinal neurons in cleared brain tissue. Whole brains were cleared using 3DISCO and imaged by LSFM wo weeks after lumbar injection of AAV2-retro-mScarlet or H2BmScarlet. (a,b) Dorsal views of whole brains with cytoplasmic (a) and nuclear-localized (B) mScarlet. (C, D) High magnification views of cerebral cortex showing corticospinal tract neurons with cytoplasmic or nuclear label. (e-h) Dorsal view of midbrain (E,G) and brainstem (F,H). Nuclear localized signal in G,H shows no interference from axon tracts and increases the number of visible cells (arrows). (i) Higher magnification of cerebral cortex neurons labeled with nuclear localized mScarlet. (J) Yellow dots corresponding to the labeled nuclei in I. (k) Merged I and J images. I, Quantification of cells expressing the cytoplasmic and nuclear localized mScarlet in cerebral cortex , red nucleus and brainstem. Scale bar, a,b, 1000 µm; c,d, 200 µm; e-h, 500 µm. OB, Olfactory bulb; PT, Pyramidal tract; BP, basilar pons; SC, spinal cord; CST, cortico spinal tract; RN, Red nucleus; PNN/PB, Pedunculopontine and Parabrachial nuclei. **p, .01, 2-Way Anova with post-hoc Sikak’s, N = 4 animals per group.

We next tested a strategy of localizing mSc signal to cell nuclei by histone 2B (H2B) fused in frame. Compared to cytoplasmic expression, nuclear localization appeared to greatly increase the number of discernable objects in cleared tissue (**Figure 1 E-H).** Indeed, spot detection confirmed an approximately two-fold increase in the number of detected cells in the cortex, and more than 10-fold increase in the brainstem (249.6 ± 56.3 SEM vs. 3334.3 ± 82.8 SEM, **Figure 1 I-L**). Close examination of spot detection of nuclear localized label showed that even in areas of relatively densely packed supraspinal neurons the cell nuclei were well spaced, enabling effective segregation and detection (**Figure 1I-K)**. We conclude that compared to cytoplasmic label, nuclear-localized fluorescence greatly enhances cell detection in cleared tissue and utilized H2B constructs throughout this study.

To further optimize cell detection we tested mGreenLantern (mGL), a recently described green FP with enhanced brightness ^26^. To directly compare mGL to mSc, adult mice received lumbar (L1-L2) injection of mixed AAV2-retro-H2B-mGL and -mSc, followed two weeks later by brain clearing, 3D imaging, and nuclei detection using Imaris software. Based on location we classified retrogradely labeled nuclei into six groups: corticospinal, hypothalamic, red nucleus, dorsal pons, medullary reticular formation, and caudal dorsal medulla (**Figure 1 – figure supplement 1A-N**). Note that additional nuclei existed outside these easily recognizable areas and are considered below. Quantification of labeled objects revealed that H2B-mGL significantly increased detection of neurons in cortex, dorsal pons, and reticular formation (p<.01, 2-Way ANOVA with post-hoc Sidak’s) (**Figure 1 – figure supplement 1L-O**). Counts were similar in right and left hemispheres, confirming that spinal injections reached both sides equally (**Figure 1 – figure supplement 2A**). Counts also did not increase between two and four weeks post-injection (**Figure 1 – figure supplement 2B**). Combined, these data establish an initial categorization and quantification of supraspinal brain regions in 3D space, reveal H2B-mGL to be the most sensitive of the FPs tested, and create consistent experimental parameters for the detection of supraspinal neurons.

### A pipeline for detection and spatial registration of supraspinal projection neurons

The procedures outlined above improve detection of supraspinal nuclei yet remain reliant on user-generated judgements regarding cell location. This is likely only practical and accurate for large and isolated populations, and new approaches are needed for brain regions with closely adjacent or intermingled populations ^27^. We therefore established an analysis pipeline to standardize brain registration and cell detection, using tools and concepts derived from the Brainglobe initiative (**Figure 2A-D**). After tissue clearing and initial inspection in Imaris, image stacks were exported and pre-processed to create cell and background sets in standard orientation (**Figure 2A,B**). Images were then registered and segmented using automated mouse atlas propagation (aMAP/brainreg), a well-validated tool to align 2D datasets with the 25um version of Allen Mouse Brain atlas ^20, 21, 28^ (**Figure 2C**). We next used cellfinder, a deep learning model-based tool, for identification of labelled cells in whole brain images ^21^ (**Figure 2C**). In conjunction with aMAP/brainreg, cellfinder assigns objects to 645 individual brain regions and quantifies the number in each. In addition, cellfinder produces detailed visualization of each optical slice, with defined brain regions outlined and labelled cells represented as overlaid spots (**Figure 2C, Figure 2 – figure supplement 1**). For final visualization we used Brainrender ^29^ which displays cellfinder output in an interactive 3D format registered to the Allen Mouse Reference Atlas (**Figure 2D**). To disseminate the insights from this pipeline, both 2D and 3D visualizations of the brains described below are available on an interactive web interface (3Dmousebrain.com).

**Figure 2:**
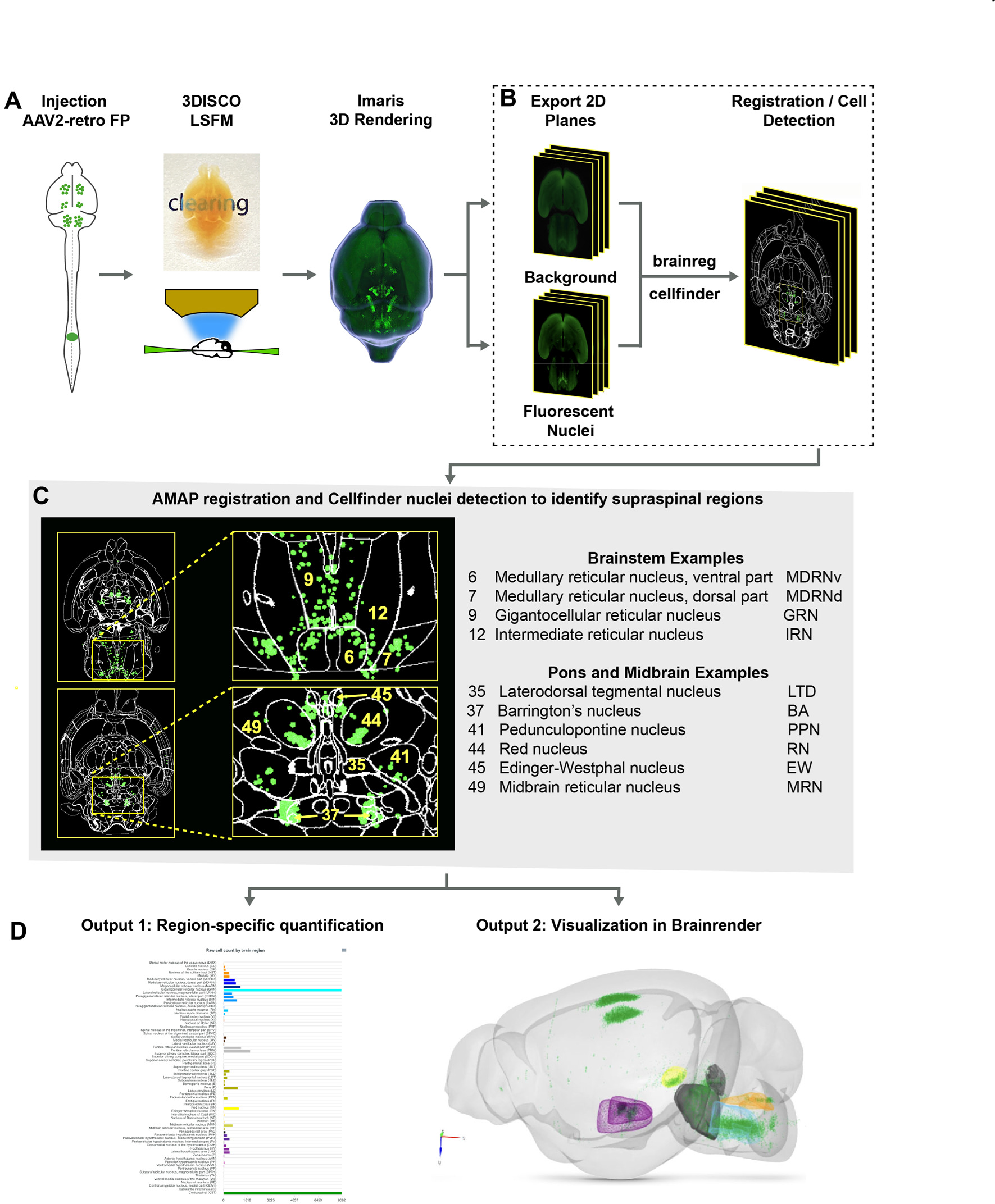
A pipeline and web resource for the detection and spatial registration of supraspinal projection neurons. **(A)** Tissue preparation and initial imaging. AAV2-retro-H2B­ mGL is injected to the spinal cord, followed by perfusion, tissue clearing, light-sheet imaging, and 30 processing using lmaris software. **(B)** Image registration. A complete series of background and fluorescent nuclei images are exported for registration to standard 30 space by brainreg and cell nuclei detection by cellfinder. (C) An example of cellfinder output, showing horizontal brain sections with brain regions outlined and detected cell nuclei indicated in green. **(D)** Example output available on “https://30mousebrain.com.” On the left are quantitative nuclei counts for identified brain regions, and on the right is 30 visualization of supraspinal locations generated by Brainrender, an interactive python-based tool.

### Brain clearing and registration quantifies supraspinal connectivity to the lower spinal cord

We first applied the registration pipeline to examine connectivity from the brain to the lower spinal cord. Ten animals received injection of AAV2-retro-H2B-mGL to L1 spinal cord, followed two weeks later by perfusion, imaging, and analysis. On average, 31,219 nuclei were detected per brain (range 20,688 to 40,171). Complete nuclei counts are available in **Source Data 1**. Below we highlight the variety of supraspinal populations identified by 3D registration, with reference to prior descriptions in rodent that help validate the automated findings. Note that we adopt nomenclature from the Allen Mouse Reference Atlas, which the cellfinder pipeline employs.

In the medulla the largest group of supraspinal neurons registered to the gigantocellular nucleus (GRN), an evolutionarily conserved source of both pre-autonomic and pre-motor axons (**Figure 2 – figure supplement 1B.8**) ^30–33^. More ventrally, labeled nuclei were present in the magnocellular reticular nucleus (MARN), which also project to the ventral horn and IML of the lower spinal cord (**Figure 2 – figure supplement 1B.10**). Notably, labeled nuclei also mapped to regions lateral to the GRN, including the Paragigantocellular Reticular Nucleus, lateral part (PGRNl) (**Figure 2 – figure supplement 1B.10**). This region contains spinally projecting neurons that initiate locomotion, as well as the ventral rostral medullary group that regulates blood pressure ^34, 35^. A cluster of labeled nuclei was also located dorsally in the caudal medulla, within and near the solitary nucleus, as supraspinal region implicated in visceral input to respiration and cardiovascular tone ^7, 8, 36^ (**Figure 2 – figure supplement 1.3**).

More rostrally in the brainstem, labeled nuclei were present in the spinal and medial vestibular nuclei, which project to spinal targets to mediate postural control (**Figure 2 – figure supplement 1B.3**). Labeled nuclei were also abundant in the pontine reticular nuclei ^7, 8^ (**Figure 2 – figure supplement 1B.6**). Although perhaps less well understood than medullary reticular populations, pontine reticular neurons have been linked to muscle atonia during sleep, startle responses, and to multi-segment postural adjustments during limb extension ^37, 38^. More dorsally in the pons, labeled nuclei mapped to known supraspinal regions in and around the pontine central grey, including the locus coeruleus (LC), laterodorsal and sublaterodorsal tegmental nucleus ^7, 8, 39– 41^ (**Figure 2 – figure supplement 1B.3**). Labeled nuclei also registered to Barrington’s nucleus (BAR), which plays a central role in the control of micturition and bowel control ^42, 43^ (**Figure 2 – figure supplement 1B.2**). Another prominent nucleus was the parabrachial nucleus (PBN), a sensory relay for inputs related to itch, pain, touch, and a range of autonomic controls including blood pressure and thermoregulation (**Figure 2 – figure supplement 1B.3**). (Chiang et al, 2019, Choi et al, 2020).

In the midbrain, supraspinal nuclei were prominently detected in the red nucleus (RN), as expected, and in midline regions including Edinger Westphal (EW) and the Interstitial Nucleus of Cajal (INC), whose supraspinal projects are known to affect postural adjustments and energy homeostasis (**Figure 2 – figure supplement 1B.2**) ^7, 44, 45^. Interestingly, nuclei also mapped to the midbrain reticular nucleus and pedunculopontine nucleus (PPN) (**Figure 2 – figure supplement 1B.2**), which lie within the mesencephalic locomotor region (MLR), a well-studied region in which stimulation triggers locomotion in a range of species including mice ^46, 47^. Although much MLR activity acts through reticular relays, the presence of direct supraspinal input from the PPN and mesencephalic reticular nucleus (MRN), noted here and elsewhere, indicate some role for direct spinal activation ^46, 48–50^.

Finally, in the forebrain, large clusters of nuclei were detected in the corticospinal tract region, as expected. These are considered in more detail below. Supraspinal neurons were also detected in the hypothalamus, where they separated into two prominent clusters, medial and lateral (**Figure 2 figure supplement 1B.7**). The medial cluster mapped mostly to the paraventricular hypothalamic nucleus (PVH) and the adjacent descending paraventricular nucleus (PVHd). These are known to innervate autonomic circuitry in the lower spinal cord and to modulate functions including bladder control, sexual function, and blood pressure ^51–53^. The lateral cluster spanned the dorsalmedial nucleus (DMH) and the lateral hypothalamic area (LHA). Although less well characterized than the PVH, prior work in the LHA has identified orexin-expressing neurons that project to all spinal levels with functions that include pain modulation ^54, 55^.

Overall, 3D imaging and registration located tens of thousands of neurons across the neuroaxis. Importantly, supraspinal neurons were mapped to distributed regions with broad correspondence to existing understanding of supraspinal connectivity. We also tested for variation in supraspinal numbers when injection sites were adjusted slightly to either lower lumbar (L4) or lower thoracic (T10). Interestingly, compared to L1 these cohorts showed no significant differences beyond a modest increase in CST and a modest decrease in GRN in T10-injected animals, highlighting the consistency of the approach (**Figure 3**). We conclude that 3D imaging and neuro-anatomical registration provides a quantitative and global profile of neurons with spinal projections.

**Figure 3:**
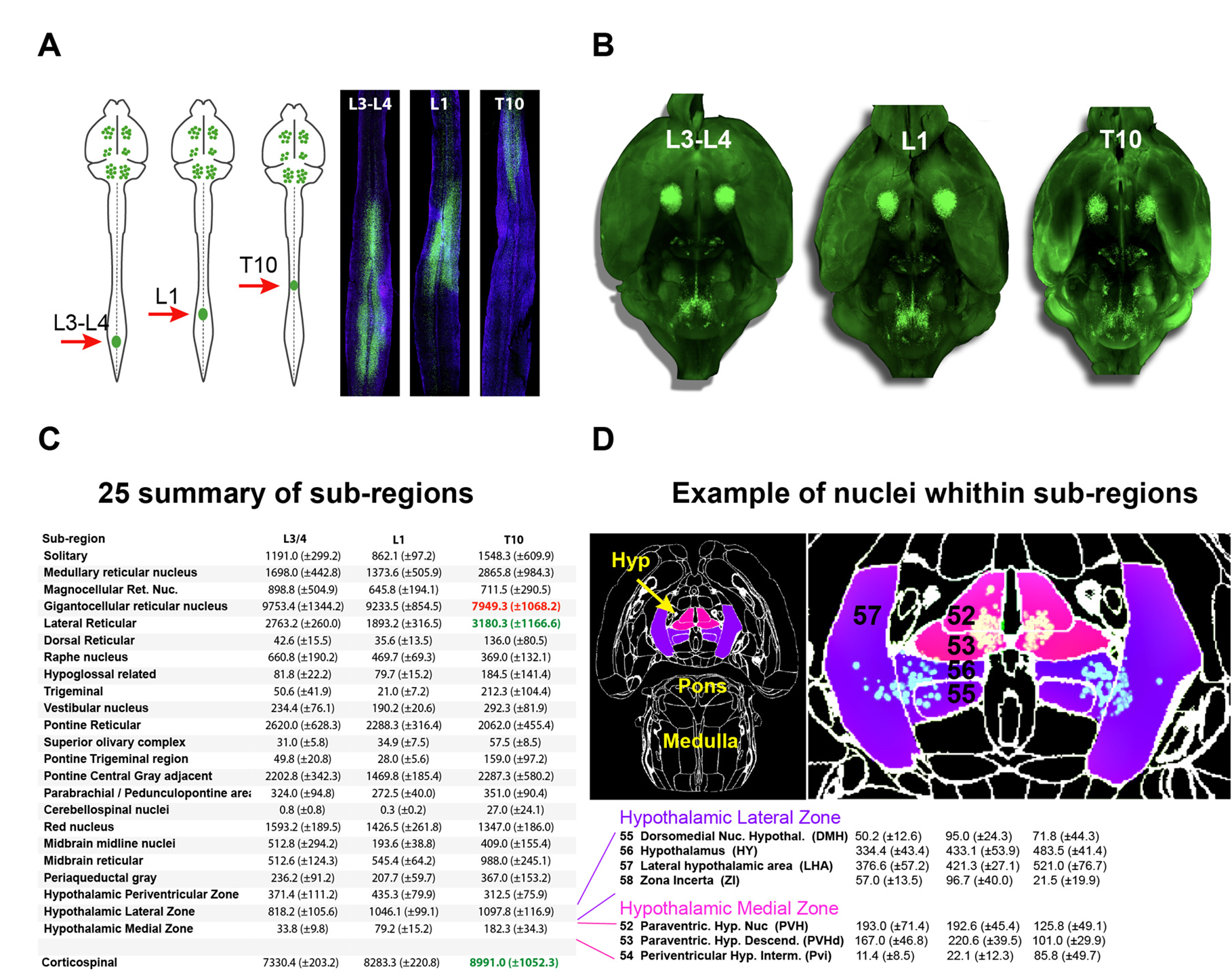
Registration of mouse brains injected at T10 , L1 and L3-4 show a similar pattern of labelled supraspinal neurons. (a) AAV2 -retro-H2B-mGL was injected into the SC at the indicated levels (red arrows). Two weeks later animals were sacrificed and horizontal sections of spina l cord prepared, with green nuclear-localized label confirming correct targeting of injections. ( b) 30 Ventral views of cleared and imaged brains of the same animals as in A , showing qualitatively similar distribution of supraspina l neurons (green label). (c ) Quantitative assessment of defined supraspinal regions using cellfinder shows similar counts in all but three regions. Green text marks significant elevation and red significant reduction (p<0.05 , 2-WAY ANOVA with TUKEY’s Multiple Compariso n, N=4 anima ls) (d) An examp le of nuclei localization and counts within in sub-regions of the brain using aMAP and cellfinder, highlighting the hypothalamic lateral zone and medial zone.

### Use case 1: Brain-wide comparison of supra-lumbar versus supra-cervical connectomes

Supraspinal populations can display topographic mapping with respect to spinal levels. For example, motor cortex is divided loosely into forelimb and hindlimb regions and in the red nucleus lumbar-projecting neurons reside ventral/medial to cervical-projecting ^56, 57^. For many supraspinal inputs, however, little is known regarding potential differences. We therefore performed 3D imaging and registration in animals that received cervical injection of AAV2-retro-H2B-mSc and lumbar injection AAV-retro-H2B-mGL, enabling within-animal comparison (**Figure 4A-M)**. We first examined overall differences in cell counts between mSc (cervical) and mGL (lumbar) label across brain regions by calculating the percent of total labeled nuclei that contained mSc signal, creating an index of cervical preference. Note that because cervical label was more abundant, comprising an average of 73% of nuclei throughout the brain (**Figure 4N**), we considered this level to be baseline and focused on deviation from it in individual areas.

**Figure 4:**
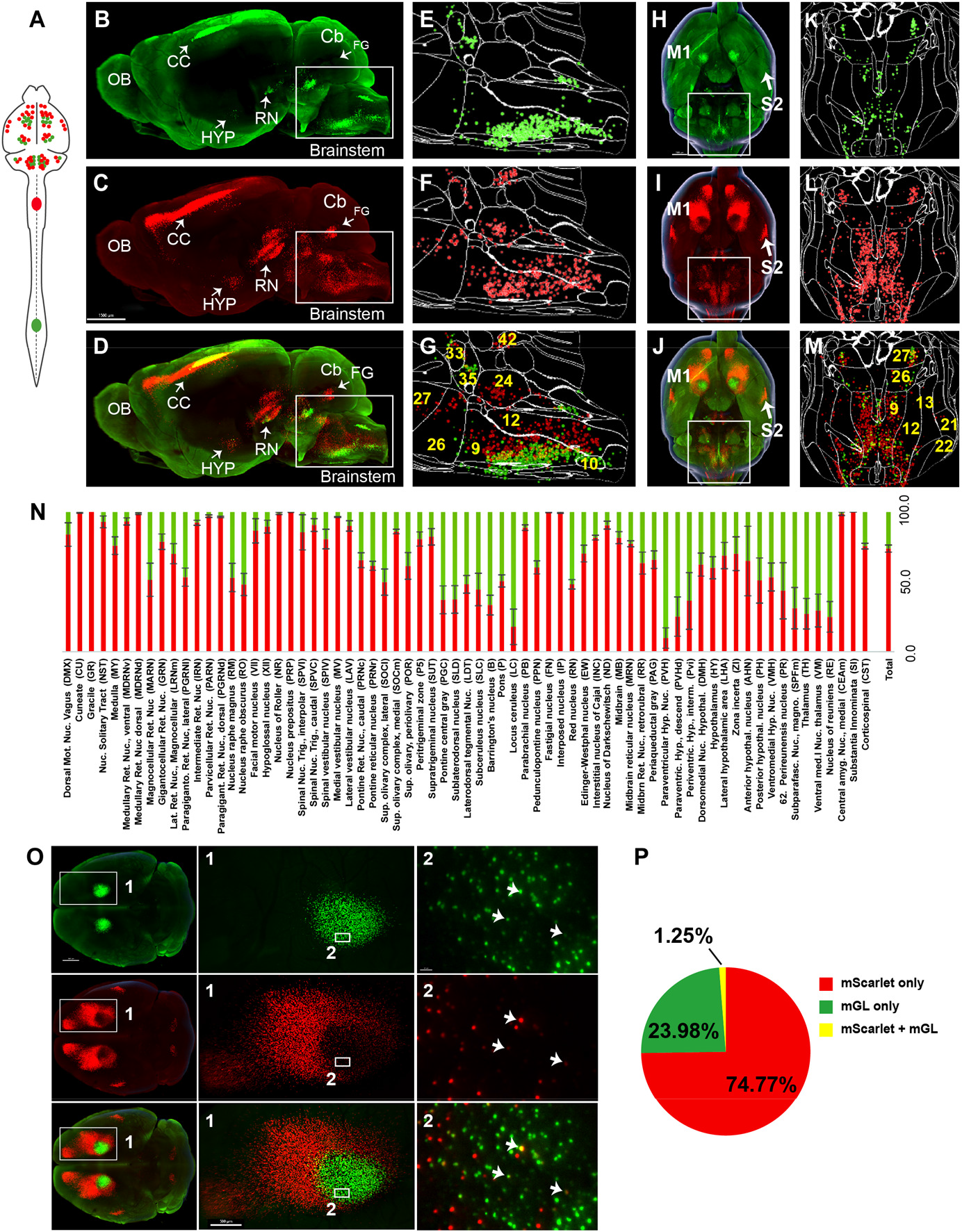
A brain-wide quantitative comparison of cervical and lumbar-projecting supraspinal neurons. (A) Experimental approach. AAV2-retro-H2B-mSc and mGL were delivered to C4 and L1 SC respectively, followed four weeks later by brain clearing, light-sheet microscopy, registration, and quantification. (B-G) Lateral view of brain and cellfinder output from brainstem regions. b,e show mGL (lumbar), C,F shows mSc (cervical) and D,G show the overlay. Note the greater abundance of cervical signal in dorsal brainstem. (H-M) Dorsal brain views and cellfinder output of the same animals as b-g. h,h show mGL (lumbar), 1 ,L show mSc (cervical), and J,M show the overlay. Note the relative abundance of cervical label in more lateral brainstem. (N) Quantification of the percent of cells in each brain region that project to cervical (red) or lumbar (green) spinal cord. (0) 30 dorsal views of cortical neurons labelled in green from L1 and red in C4. Inset 1 shows the entire M1 region, inset 2 shows intermingled cervical and lumbar region. Most cell nuclei are single-labeled; arrows show exceptions. (P) Quantification of the percent of neurons in M1 labeled with only mGL, only mSc, or dual-labeled. Scale bar, B,C,D, 1500 µm; H,l,J, 1000 µm; o, Left pictures 1000 µm, middle 500 µm, right 30 µm. n = 4 biological replicates per group. OB, Olfactory bulb; CC, Cerebral Cortex; M1, Motor area 1; HYP, hypothalamus; RN, red nucleus; Cb, Cerebellum; Fastigial nucleus; P, Pons; M. medulla.

Several patterns emerged. The medullary reticular formations, which occupy the most caudal region of the medulla, and the laterally positioned parvicellular nucleus both showed a predominant cervical projection. Interestingly, these regions have been recently implicated in forelimb reaching behavior, a function that would be consistent with predominantly cervical projections ^58, 59^. The 3D registration approach also revealed an overall topographic pattern in the brainstem in which ventrally located populations projected to both lumbar and cervical regions, whereas more dorsally located populations were predominantly cervical (**Figure 4B-G**). The nucleus prepositus and Roller nucleus, dorsally located in the medulla, showed mostly cervical projections, possibly related to their known role in gaze tracking (**Figure 4N**) ^60^. In contrast, the pontine reticular formations showed a more balanced distribution. The red nucleus was relatively balanced, but showed a topographic pattern in which ventral-medial neurons projected to lumbar cord (**Figure 4B-D**) ^19, 56, 61^. Neurons near the pontine central grey, including Barrington’s nucleus, showed relative enrichment for lumbar labeling, consistent with known innervation of lumbar circuitry (**Figure 4E-G**)^43^. Similarly, the paraventricular region of the hypothalamus was enriched for lumbar label, consistent with its known innervation of the IML cell column. Overall, these data use comprehensive, brain-wide approach to both quantify previously noted topographic differences in spinal level targeting and to reveal new patterns.

Next, focusing on the corticospinal tract, we used higher resolution imaging to examine within-cell colocalization of label from both cervical and lumbar cord. As expected, cervical mSc label was apparent in three discrete cortical areas: a large central mass (M1/S1), a more rostrally located cluster (Rostral Forelimb Area, or RFA), and a lateral mass (S2) (**Figure 4O**). The mGL label from lumbar cord was concentrated in a subregion of M1, centered medial and caudal to the main mass of cervically-labeled neurons. Interestingly, a rim of cervically labeled cells completely surrounded the lumbar region (**Figure 4O, Figure 4 – figure supplement 1**). Thus lumbar-projecting CST neurons can be described most accurately not as a separate caudal population, but rather as nested within a broader region of cervical-projecting neurons ^22^. In addition, even in cortical areas where cervical and lumbar label overlapped, definitive within-cell colocalization was detected in only 1.25% of neurons (**Figure 4P**). Importantly, in prior experiments when AAV2-retro-mSc and -mGl were co-injected to lumbar spinal cord, more than 80% of CST neurons expressed both fluorophores (**Figure 1B, C**), arguing strongly against any viral interference mechanism that prevents dual transgene expression. Thus, the low rate of co-expression when the two viruses are delivered separately to cervical and lumbar spinal cord indicates largely distinct innervation by individual CST neurons to only one of the two regions. Notably, a recent manuscript using a similar viral approach showed consistent, albeit qualitative, findings ^22^. Thus, although dual cervical/lumbar collateralization by single CST axons has been reported in juvenile mice ^62^, in adult mice CST collateralization is highly region-specific ^22^. In summary, 3D imaging and registration can determine both brain-wide and within-population patterns of topographic innervation across spinal levels.

### Use case 2: Application to spinal injury

We next applied whole-brain imaging and quantification to questions related to spinal cord injury. First, in light of the large existing patient population and the practical difficulties in delivering therapeutics immediately after injuries, a critical question for the eventual clinical use of gene therapy vectors is their capacity for effective transduction when applied in conditions of chronic injury. Our prior work showed AAV2-retro’s efficacy in some cell types immediately after injury^19^, but this conclusion was only qualitative, applied to limited supraspinal populations, and did not examine more extended and clinically relevant time points. We therefore injected AAV2- retro-H2B-mGL six weeks after a complete crush of thoracic spinal cord (**Figure 5A**). After sacrifice, examination of the crush site in sections of spinal cord confirmed injury completeness, as evidenced by a lack of astrocytic bridges and lack of retrograde label distal to the injury (**Figure 5B**). In cleared brains, examination of retrograde mGL showed a broad distribution of signal (**Figure 5C-G**), and in all regions the nuclei counts did not differ significantly from those found previously in uninjured animals with similar thoracic injections (**Figure 5H**). These data quantitatively verify AAV2-retro’s ability to effectively deliver transgenes to a wide diversity of cell types in the chronic phase of injury, meeting an important criteria for eventual translation.

**Figure 5.**
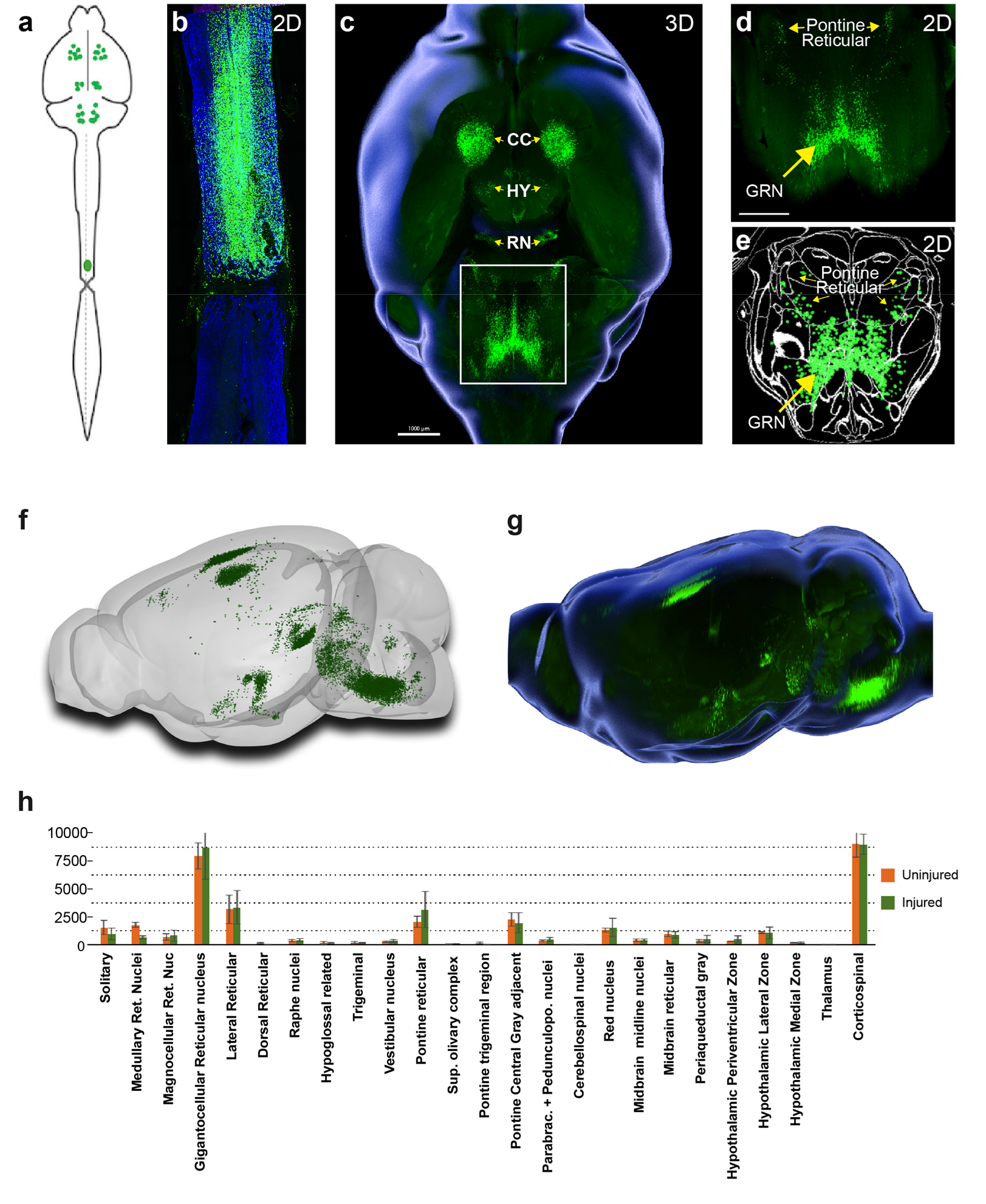
Retro-AAV2 effectively transduces neurons when delivered to the chronically injured spinal cord. (a) Experimental design. A complete crush injury was delivered to lower thoracic spinal cord, followed six weeks later by injection of AAV2-retro-H2B-mGL rostral to the injury. Two weeks post­ injection animals were euthanized, brains were cleared and imaged with light-sheet microscopy , and images were processed for registration and quantification by aMAP and cellfinder. **(b)** Horizontal spinal section stained for GFAP (blue), confirming the complete crush and verifying transduction (green) rostral but not caudal to the injury. (c-e) Ventral views of the brain from the same animal of section in **b.** C shows a 30 overview, 0 shows a higher magnification view of the brainstem, and **E** shows cellfinder output with retrograde mGL detection in green. **(f)** Brainrender output showing lateral views of the same brain in C. **(g)** lmaris 30 generated equivalent lateral view of the brain in **F. (h)** Quantification of retrograde nuclei detected in 25 brain regions, comparing uninjured animals (orange) to chronically injured animals (green). No regions displayed statistical differences (p>.05, 2-WAY ANOVA with post-hoc Sidak’s). N = 4 animals per group.

We next applied whole-brain quantification to a central challenge in SCI research, the issue of injury variability. In both the clinic and the laboratory, spinal injuries are often incomplete and leave inconsistent numbers of spared connections in each individual ^63^. To generate a range of injury severities adult mice received crush injury to T10 spinal cord using stoppers of defined thickness to produce mild, moderate, or severe damage ^64^. AAV2-retro-H2B-mGL was injected to L4 spinal cord seven weeks post-injury and tissue was analyzed two weeks post-injection. First, viral targeting and injury severity were assessed in spinal tissue sections (**Figure 6A-F**; images of all spinal cords are provided in **Figure 6 – figure supplement 1**). As expected, mild injuries displayed elevated GFAP at the crush site but overall astrocytic continuity (**Figure 6A, B**). In contrast, severe injuries showed complete gaps in GFAP at the site of injury, with no detectable astrocytic bridges (**Figure 6E,F**). Moderate injuries were highly variable and sometimes asymmetrical right to left (**Figure 6C, D**). Overall, as intended, the injuries displayed a wide range of severity.

**Figure 6:**
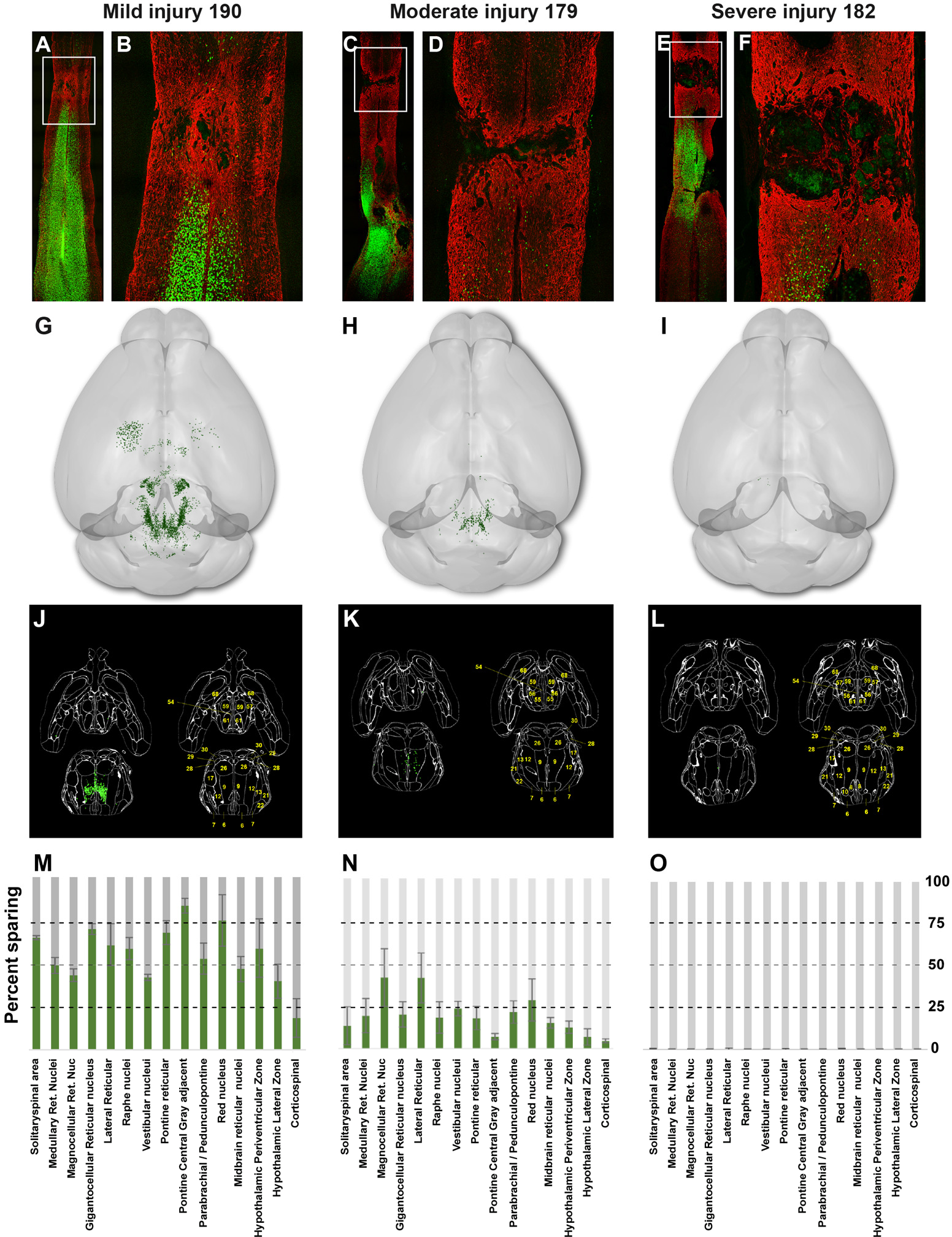
Brain clearing and registration quantifies region-specific sparing across a range of spinal injury severities. (A-F) Animals received thoracic crush injuries of controlled widths, followed eight weeks later by injection of AAV2-retro-H2B-mGL to L1 spinal cord. Horizontal spinal cord sections show mild (A,B), moderate (C,D), or severe (E,F) injury, with GFAP in red and viral transduction in green. (G-L) Dorsal views of the brains from the same animals of the spinal cord sections above, showing progressive reduction in the number of retrogradely labeled neurons as injury severity increases. G-1 show Brainrender depictions of whole brain, J-L show one 20 cellfinder plane output with brain regions outlined and detected mGL in green on the right panels. (M-0) Quantification of percent sparing in identified brain regions, with the average value from uninjured animals set as 100. N = 3 mild, 6 moderate, and 3 severely injured animals.

To assess residual brain-spinal cord connectivity supraspinal neurons were registered and quantified by the pipeline described above (**Figure 6G-L**). Raw values for all brain regions are provided in **Source Data 1,** and **Figure 6M-O** shows values normalized to region counts in uninjured mice, thus creating an index of sparing for each region. Mice that received severe injuries showed a maximum of 29 labeled cells brain-wide, confirming disruption of descending axon tracts (**Figure 6O**). In contrast, mild injuries averaged only a 43% reduction in retrograde label, with high variability between different supraspinal populations (**Figure 6J**,**M**). For example, the CST was strongly affected, averaging less than 20% sparing, while neurons near the pontine central grey and the red nucleus averaged 83.3% and 75.1% sparing, respectively (**Figure 6M**). The moderate injury group showed a brain-wide average of 24.2% sparing, also with high variability between animals (range 3.4% to 36.3%) (**Figure 6K,N**). As in the mildly injured animals, CST neurons were affected more strongly than other populations such as the PGC and RN, although the animal with the highest overall sparing showed an unusual pattern of nearly 70% persistence of the corticospinal tract (**Figure 6N**). In summary these data confirm the ability of brain-wide analysis to detect overall sparing differences in groups of animals that received injuries of different severity, and more importantly to detail differences in the injuries’ effects on individual animals and on individual cell populations.

We next asked how indexes of sparing correlate with functional recovery from spinal injury as assessed by the BMS motor score, a well-established measurement of hindlimb function and interlimb coordination ^65^. As expected, BMS scores averaged lower in animals that received severe injury versus moderate or mild (**Figure 7 – figure supplement 1**). Notably, however, within each group the scores varied widely. Combining all injury groups, we first examined the correlation between BMS scores and the size of the spinal injury, measured in spinal sections and defined as the GFAP-negative region bounded by gliosis. We identified a significant negative correlation between injury size and BMS score, but consistent with many prior findings in the field, injury size could explain only some of the variability in the data (R^2^=0.53; **Figure 7 – figure supplement 1B-E**).

**Figure 7:**
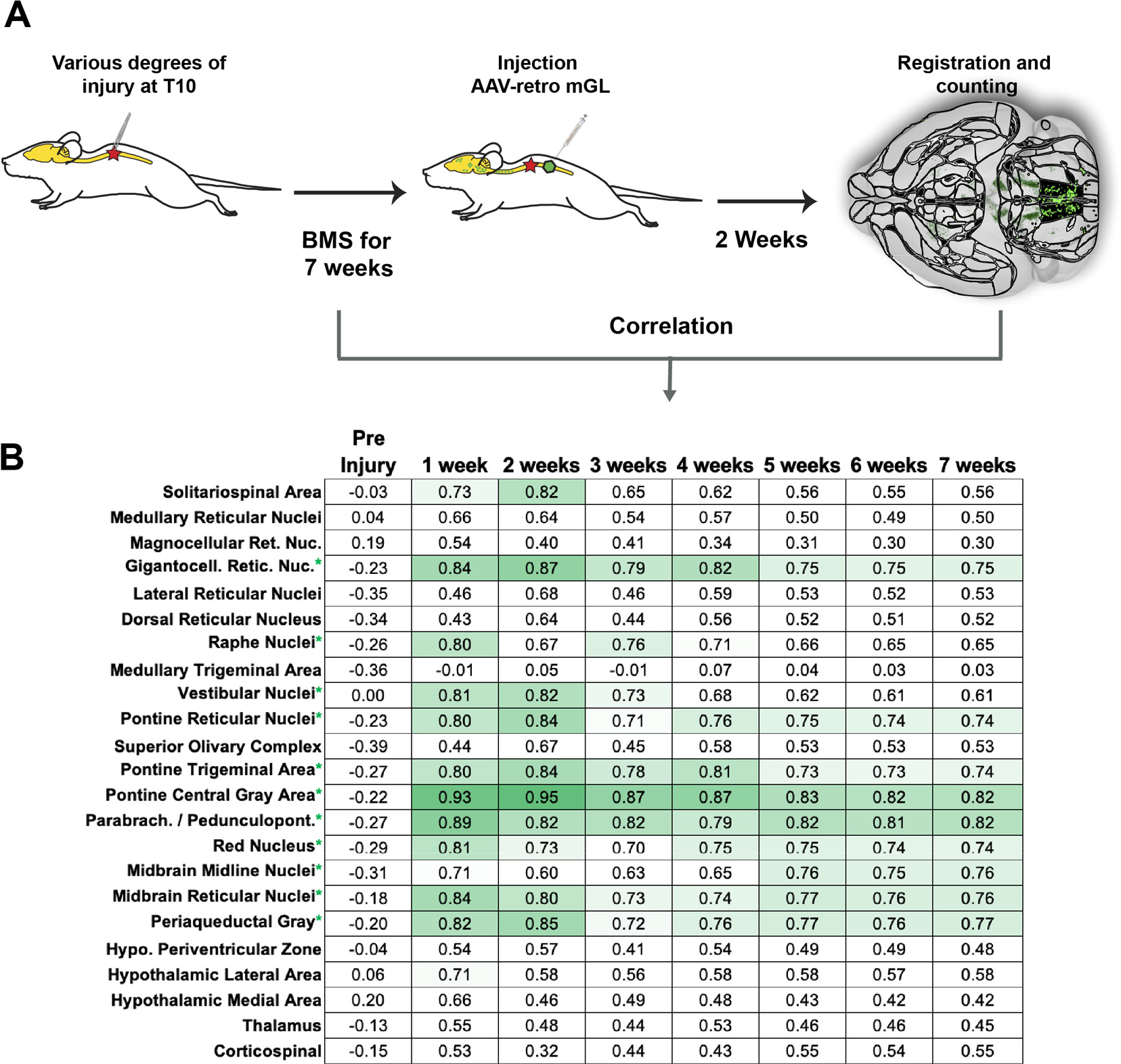
Residual connectivity in specific brain regions correlates with motor recovery after spinal cord injury. (A) Experimental design. Adult mice received graded spinal cord injuries followed by weekly BMS testing. Seven weeks post-injury animals received lumbar injection of retro-AAV2-H2B-mGL , followed two weeks later by perfusion, 30 imaging of brains, registration, and region-specific quantification of spared neurons. (B) shows a correlation matrix of R values derived from linear regression between the number of spared neurons in each of 25 brain regions and weekly BMS scores. * indicates eleven regions the regression slope differed significantly from zero (p<.05, simple linear regression)

We hypothesized that variability in motor recovery may also reflect inter-animal differences in the amount of sparing of supraspinal neurons in selected brain regions. We therefore created a correlation matrix to test for the potential relationship between locomotor scores and the number of spared neurons in each individual brain region (**Figure 7A,B).** We made the assumption that because minimal long-distance regeneration occurs spontaneously, the spared neurons detected at the terminal 7-week timepoint likely matched those in prior weeks, and therefore tested for correlation between spared neurons and BMS scores at each time point. Counts in many regions, such as the CST, hypothalamus, and lateral reticular formation correlated poorly with hindlimb function, and regression slopes did not differ significantly from zero (R<0.5, p>.05, simple linear regression). In contrast, in ten brain regions the regression slopes differed significantly from zero, with R values between 0.75 and 0.93 at different time points. Included in these were regions relatively well studied in locomotor recovery after spinal injury, notably the red nucleus and gigantocellular reticular. Interestingly, other correlated regions, for example the pedunculopontine and periaqueductal grey, have received less attention in spinal injury research but have been functionally linked to locomotion ^46, 47^. In addition the number of spared propriospinal neurons in cervical spinal cord, counted in cleared tissue between C2 and C6, also correlated well with functional recovery, highlighting the potential importance of cervical neurons as a supralumbar control center (R=0.83, slope<0 p=.0018, simple linear regression) (**Figure 7 – figure supplement 2**) ^66^. Overall, these data support the utility of whole-brain imaging and quantification to partially explain variability in functional outcomes after spinal injury.

## Discussion

We present a novel experimental approach and associated web-based resource that provides comprehensive quantification and visualization of neurons that project from the murine brain to specific levels of the spinal cord. Supraspinal projecting populations are numerous and diverse, yet prior publications, particularly in the spinal cord injury field, have focused disproportionately on specific sets of nuclei and a handful of major pathways. The new approach presented here is needed to spur progress by clarifying the complex geometry of supraspinal neurons and offering a practical means to assess post-injury connectomes across the entire brain in numerous animals within an experimental study. This approach opens the door to comprehensive analyses of changes in supraspinal connectivity in response to disease and injury, and conversely to profile without bias the brain-wide efficacy of pro-regenerative therapeutics.

### Improved cell detection in optically cleared tissue

An essential element of this approach is the deployment of newer-generation FPs, mScarlet and mGreenLantern, with nuclear targeting ^25, 26^. Compared to prior vectors we showed strongly enhanced detection of retrogradely transduced neurons, most notably in the brainstem, an important source of supraspinal control. This likely reflects the relative concentration of fluorescence in the nucleus, augmented by slower protein turnover and reduced interference from intervening axon tracts. It is likely that prior work, including our own and a recent description of CST neurons in cleared brains, underestimated the number of retrogradely transduced neurons ^19, 22^. Thus nuclear-localized mSc and mGL expand the tool kit for neuronal labeling in cleared tissue and allow flexibility in dual labeling experiments.

### Toward a more accessible supraspinal connectome

We designed our workflow to meet two central objectives: to quickly assign supraspinal neurons to precise locations, and to disseminate this insight in a cohesive, practical fashion ^27^. Prior work has employed retrograde tracing and manual scoring of brain sections to meticulously catalogue supraspinal populations, and in most cases this foundational work should be considered definitive ^7, 8^. Indeed, the broad concordance of these prior human-curated efforts with our automated registration approach, highlighted above in Results, provides essential validation ^1, 2, 7, 8, 56, 59, 60^. As one standout example, the detection of cervically-projecting neurons in the amygdala was initially surprising, but a literature search revealed a decades-old description of this pathway in monkeys and more recently in mice ^8, 67^. Indeed, prior description can be found for nearly all populations detected in our workflow. The disparate and often non-quantitative nature of prior work, however, present significant challenges to effective utilization. These challenges are compounded by the anatomical complexity and the difficulty of synthesizing anatomical data presented in 2D or described in reference to anatomical landmarks. The pipeline presented here marks an important step in overcoming these challenges by offering standardized detection, registration, and quantification of supraspinal populations across the brain. In addition, the ability to visualize populations in 3D lowers conceptual barriers by generating intuitive insights between the data and brain regions. By focusing on connectivity between the brain and spinal cord this novel resource fills a gap in existing web-based neuro-anatomical atlases, which focus mostly on intra-brain circuitry. Illustrating the utility, below we highlight several key insights of high significance to both basic neuroanatomy and to preclinical research that were enabled by the new approach.

### Global assessment of cervical / lumbar topographic mapping from supraspinal neurons

A long-standing interest in neuroanatomy is the degree to which supraspinal populations innervate spinal targets in cervical regions, lumbar regions, or both. That understanding is fundamental to interpreting supraspinal function. Our findings strongly indicate the AAV2-retro is taken up by axon terminals but minimally or not at all by fibers of passage. For example, by comparing our labeling to the known cervical/lumbar patterns withing CST and red nucleus, it is clear that AAV2-retro injected to cervical spinal cord is taken up by axons that innervate the injection site but not by nearby axons that travel to lumbar regions. Thus AAV2-retro provides a tool to establish patterns of axon termination. One important finding is the very low incidence of dual innervation by CST neurons of both cervical and lumbar spinal cord. Although prior work in juvenile rodents detected higher levels of dual innervation, our quantitation of single-region targeting was highly consistent across animals and agrees with recent observations in adult mice^22^. Thus, individual CST neurons in adult mice largely target cervical or lumbar cord, but rarely both. In addition, 3D imaging highlights a topographic CST organization in which an oval-shaped region of lumbar-projecting neurons is nested within a larger cervical population, as opposed to a position entirely caudal. This interesting pattern may have implications for the developmental mechanisms by which cervical versus lumbar fates are specified, for example hinting at a point source of a diffusible specifying signal at the center of the lumbar region. More broadly, brain-wide mapping revealed a topographic pattern within the brainstem such that lumbar-projecting neurons are concentrated in the ventral regions, whereas more dorsal regions are contain almost exclusively cervically-projecting axons. Another example of highly selective innervation is found in the hypothalamus, where the paraventricular region showed a strong lumbar bias. These data provide foundational neuroanatomical insight into brain-spinal connectivity.

### Retrograde gene delivery to chronically injured supraspinal neurons

Our findings have important implications for research that aims to address the needs of individuals that suffer from supraspinal disruptions such as spinal cord injury (SCI). Besides the well-studied locomotor and fine motor deficits, SCI also affects pain sensation, bladder and bowel control, sexual function, basic postural control, cardiovascular tone, thermoregulation, and even metabolism ^14^. Supraspinal populations that serve these functions are known, yet their response to injury and to attempted pro-regenerative strategies are largely uncharacterized (but see Adler et al. 2017 for an example of CNS-wide profiling) ^68^. Treatments can advance even to clinical trials with limited information on how or if they influence axon growth in tracts beyond the major motor pathways. Thus, although much progress in this direction has been made, there arguably remains a mismatch between the varied concerns of individuals suffering from SCI and the narrower anatomical focus of SCI research. In this context, the brain-wide workflow presented here offers a practical means to expand the study of non-motor systems, and to populations that likely modulate the major motor pathways. Specifically, as pro-regenerative treatments are tested in preclinical mouse models, the whole-brain pipeline presented allows the response of diverse cell types to be monitored with the throughput needed for SCI studies. As a first illustration of this strategy, we used the pipeline to determine how the chronic injury state impacts transduction by retrograde vectors. This is an important consideration for the translational prospects of gene therapy approaches to treat CNS damage. We showed quantitatively that gene delivery remained highly effective and widespread in chronic spinal injury, information that is needed to proceed with gene therapy-based treatments of the broad concerns discussed above. In summary, adoption of this connectome-level approach has the potential to sharpen the predictive power of pre-clinical work and help to better align it with the concerns of individuals with SCI.

### Implications for the neuroanatomical-functional paradox in spinal injury research

Another central challenge in the SCI field is the so-called neuroanatomical-functional paradox, which refers to the fact that the size of lesions in the spinal cord are poorly predictive of functional outcomes ^63^. This unpredictability of SCI outcomes is a major stumbling block that has likely contributed to challenges of reproducibility in the field ^69^. The recovery of various functions after spinal injury is almost certainly impacted by the amount of residual supraspinal connectivity from specific brain regions, yet the field has lacked a practical means to quantify this key variable from most supraspinal populations. We now demonstrate comprehensive quantification of the variability of residual brain-spinal connectivity across injury types and between animals. Importantly, we found lesion size per se to correlate only partially with locomotor recovery, whereas the number of spared neurons in selected supraspinal populations correlated more strongly. It is important to note that correlation between a population’s sparing index and the degree of functional recovery does not necessarily imply functional involvement; it could, for example, be caused by axon trajectories in proximity to more functionally relevant tracts. In cases where the absolute number of supraspinal axons are low, or where prior findings contradict functional involvement, the correlations are likely epiphenomenal. The value of correlating functional recovery with residual connectivity across the brain is to rapidly generate and prioritize functional hypotheses, which must be synthesized with existing information and ultimately tested directly.

As one example, our data detect a direct projection from the pedunculopontine nucleus (PPN) and the lumbar spinal cord and a high correlation between maintenance of this pathway with locomotor function after SCI. Interestingly, the PPN lies within the mesencephalic locomotor region, an area that has been recognized for more than 50 years as an important area for locomotion ^48, 70^ but which has received limited attention in spinal cord injury research. Thus, the correlational analysis presented here raises the hypothesis that in addition to rubrospinal and reticulospinal projection, the maintenance and regeneration of PPN-spinal projections may impact locomotor recovery from spinal injury. By directing attention toward lesser-studied but potentially significant regions, the integrated approach presented here can point toward the key functional experiments needed help resolve the variability in outcomes that currently challenge the field.

Important caveats to the approach should be considered. First, automated detection of nuclei likely remains imperfect and is impacted by the image quality of LSFM, notably stretching in the Z plane. Continued improvements with more isotropic acquisition in light sheet microscopy and in trained detection of nuclei will likely resolve these lingering issues ^71, 72^. A second caveat regards viral tropism, for example AAV2-retro appears less effective at transducing serotonergic cell types, thereby limiting assessment of raphe-spinal projections ^19^. This limitation will likely be addressed as additional retrograde variants are made ^73^. Finally, it is important to note that we have applied this approach only to descending inputs to the spinal cord, and not to ascending tracts. In principle a similar retrograde strategy could quantify neurons that give rise to ascending input, and a promising future direction would be to incorporate this information into predictive models for function after partial spinal injury. However, while these and other future developments are likely to further improve the approach, the present iteration provides information on an unprecedented scale and has yielded new insights into the complexity of supraspinal populations and their variable response to spinal injury.

## Methods and Materials

### Animal information

All animal procedures were approved by the Marquette University Institutional Animal Care and Use Committee and complied with the National Institutes of Health Guide for the Care and Use of Laboratory Animals. Adult female C57BL/6 mice (6-8 weeks old, 20–22 g) were used for these experiments. Age at day of surgery was eight weeks and mean weight was 20g. Group for initial fluorophore optimization were: mScarlet cytoplasmic, L1 injected, 4 weeks: 4 animals; H2B-mScarlet, L1 injected, 4 weeks: 4 animals; H2B-mScarlet, L1 injected, 2 weeks: 8 animals; H2B-mGreenLantern, L1 injected, 2 weeks: 8 animals. Group sizes for cleared and registered brains were: L1 injected: 10 animals; L3/4 injected: 5 animals; T10 injected: 4 animals; Cervical/lumbar co-injected: 4 animals; chronically injured: 3 animals; moderately injured: 6 animals; mildly injured: 3 animals; severely injured: 3 animals. In the 2 week, L1 injected groups one H2B-mGl-injected animal showed likely blockage of the injection needle as evidenced by lack of local transduction at the injection site, and on H2B-mSc-injected animal suffered tissue damage during the brain dissection. These animals were excluded, leaving 7 per group. The room temperature was set at 22°C (±2°C) and room humidity was set at 55% (±10%).

Mice were kept in a 12-h light/dark cycle with access to food and water *ad libitum*. Mice were checked daily by animal caretakers.

### Plasmid construction and cloning

We used 2 monomeric bright fluorescent proteins (FP) of similar size that encode for mGreenLantern ^26^ and mScarlet ^25^ and fused in frame with the core histone H2B in the amino terminus for nuclear localization of the FPs. Both fusions were synthetically constructed (Genscript, USA). The AAV mGreenLantern was constructed first by generating a synthetic cDNA optimized to the Human codon usage. The rat gene H2B/Histone H2B type 1- C/E/G (accession #NP_001100822) was fused in frame to mGreenLantern with a linker of 8 amino acids (PPAGSPPA) between H2B and mGreenLantern. The fusion protein was cloned into pAAV-CAG-GFP (Addgene #37825) by substituting the GFP with H2B- mGreenLantern using restriction enzymes BamHI and XhoI. For the mScarlet the human H2B clustered histone 11 (H2BC11) (accession #NM_021058) was fused in frame without a linker and cloned into the pAAV-CAG-tdTomato (Addgene #59462) using the sites KpnI and EcoRI at the 5 and 3 prime end respectively. Cytoplasmic mScarlet was cloned identically but with the H2B sequence omitted. rAAV2-retro-H2B-mGreenLantern was produced at the University of Miami viral core facility at the Miami Project to Cure Paralysis, titer = 1.4x1013 particles/ml. Virus was concentrated and resuspended in sterile HBSS and used without further dilution. The rAAV2-retro- mScarlet and rAAV2-retro-H2B-mScarlet was made by the University of North Carolina Viral Vector Core, titer = 4.3X10^12 and 8.7x1012 particles/ml, respectively.

### Spinal cord surgery

rAAV-retro particles (1µl) was injected into the spinal cord with a Hamilton syringe driven by a Stoelting QSI pump (catalog #53311) and guided by a micromanipulator (pumping rate: 0.04 μL/min). AAV viral particles were injected at C4-C5, T10, L1, L4 vertebrae, 0.35 mm lateral to the midline, and to depths of 0.6 and 0.8 mm. For the spinal cord crush injuries adult female C57BL/6 mice (6-8 weeks old, 18–22 g) were anesthetized by ketamine/ xylazine. After laminectomy of vertebra T10-12 forceps with stoppers of defined width (0.15 or 0.4 mm) were used to laterally compress the spinal cord for 15 seconds, then flipped in orientation and reapplied at the same site for an additional 15 seconds. In one set of animals, forceps were placed laterally to the cord and within the vertebral column but not squeezed, resulting in some tissue displacement and mild injury.

### Tissue clearing and imaging

After 2-4 weeks of viral expression, animals underwent transcardial perfusion with 0.9% saline and 4% paraformaldehyde solutions in 1×-PBS (15710, Electron Microscopy Sciences). Whole brains and spinal cords were dissected and fixed overnight in 4% paraformaldehyde at 4°C and washed three times in PBS pH 7.4, followed by storage in PBS. The dura was carefully and brains and spinal cords were cleared using a modified version of the 3DISCO ^19, 74^. Samples were incubated on a shaker at room temperature in 50, 80, and twice with 100% peroxide-free tetrahydrofuran (THF; Sigma-Aldrich, 401757) for 12 hr each for a total of 2 days. Peroxides were removed from THF by using a chromatography column filled with basic activated aluminum oxide (Sigma- Aldrich, 199443) as previously described (Becker et al., 2012). On the third day, samples were transferred to BABB solution (1:2 ratio of benzyl alcohol, Sigma-Aldrich, 305197; and benzyl benzoate, Sigma-Aldrich, B6630) for at least 3 hr. After clearing, samples were imaged the same day using light-sheet microscopy (Ultramicroscope, LaVision BioTec). The ultramicroscope uses a fluorescence macro zoom microscope (Olympus MVX10) with a 2× Plan Apochromatic zoom objective (NA 0.50). Image analysis and 3D reconstructions were performed using Imaris v9.5 software (Bitplane, Oxford Instruments) after removing autofluorescence using the Imaris Background Subtraction function with the default filter width so that only broad intensity variations were eliminated.

### Imaris reconstructions

Image analysis and 3D reconstructions were performed using Imaris v9.5 software (Bitplane, Oxford Instruments) after removing autofluoresence using the Imaris Background Subtraction function with the default filter width so that only broad intensity variations were eliminated. Additionally, the entire brain was defined as an ROI in order to mask all background fluorescence outside the spinal cord surface. Artifact and nonspecific fluorescence surrounding the brain was segmented and removed using the automatic isosurface creation wizard based upon absolute intensity. Voxels contained within the created surface were set to zero and the remaining mask was used for all further analysis. Automatic segmentation of nuclei within specified ROIs was applied using the spots detection function and later superimposed on a maximum intensity projection volume rendering of the tissue. For some of the figures surfaces were created around the brains and spinal cords to make them more evident in the 3D reconstructions. Quality thresholds were set based upon visual inspection of the mixed model rendering for both spots and surfaces.

### Immunohistochemistry

Adult animals were perfused with 4% paraformaldehyde (PFA) in 1× phosphate-buffered saline (PBS) (15710-Electron Microscopy Sciences, Hatfield, PA), brains, and spinal cords removed, and post-fixed overnight in 4% PFA. Transverse sections of the spinal cord or cortex were embedded in 12% gelatin in 1× PBS (G2500-Sigma Aldrich, St.Louis, MO) and cut via Vibratome to yield 100 μm sections. Sections were incubated overnight with primary antibodies GFAP (DAKO, Z0334 1 : 500, RRID:AB_10013482), and rinsed and then incubated for 2 h with appropriate Alexa Fluor-conjugated secondary antibodies (R37117, Thermofisher, Waltham, MA, 1 : 500). Fluorescent images were acquired using Olympus IX81 or Zeiss 880LSM microscopes.

### Analysis using computational neuroanatomy

We used the BrainGlobe’s Initiative software (https://brainglobe.info) of interoperable Python-based tools for the analysis and visualization the data. For each brain we captured approximately 500 images. The captured was form ventral to dorsal to match the brainglobe default orientations (https://github.com/brainglobe). The 2D images first were assembled in Imaris v9.3.5 and 9.5 (Bitplane, Oxford Instruments). Two channels were created one to subtract the positive cell signal and generate another set with only background fluorescence. Images were exported to Image J to create a new set of TIFF files. The TIFF files of the sample images were further analyzed with a set of neuroanatomical computational tools developed for analysis of brain serial section imaging using light-sheet microscopy. First we fed, both set of images, background and positive signal images into the cellfinder a deep-learning network (Residual neural Network) to detect the positive cells (https://github.com/brainglobe/cellfinder) followed by registration and segmentation into a template brain with anatomical annotations based of the Allen Reference Mouse Brain Atlas (https://github.com/brainglobe/brainreg) and finally visualized with the brainrender (https://github.com/brainglobe/brainrender).

### Quantification and Statistics

Throughout the manuscript means are used as summary values and standard error of the mean (SEM) as the indicator of variability. Data were tested for assumptions of parametric tests using Kolmogorov-Smirnov and Levene’s tests. Values for N, the specific tests and post-hoc analyses, and p values are provided in figure legends and in the test of the Results section. All manual quantification, including behavioral assessment and measurements of lesion size, were performed by blinded observers. Statistical analyses were performed using Prism (Graphpad).

## Author contributions

P.T., and M.B. conceptualized and designed the study; Z.W., performed all the animal surgeries, viral injections, and immunohistochemical procedures; Y.W., P.T., and M.B. designed and did all the cloning of the various DNA constructs. Y.W., and P.T., cleared all the brains and image acquisition with LSFM. V.M. set up all the software tools from the Brainglobe and designed the web site. A.R. did all the analysis with Imaris. A.R., V.M., and M.B. did all the image modifications for use in cellfinder and brainrender. B.C., and G.P. designed and created the mGL. P.T. and M.B., did the analysis of all the data. M.B., wrote the first draft of the paper. B.C., P.T., and M.B., edited the manuscript. All authors contributed to the discussion of the experimental procedures, results and the manuscript.

## Acknowledgments

This work was supported by NIH/NINDS R01NS083983 (M.B and P.T.), the Bryon Riesch Paralysis Foundation (M.B.), The Miami Project to Cure Paralysis (P.T), and the Buoniconti fund (P.T). We thank Troy Margrie, Adam Tyson, Luigi Petrucco and others at the Brainglobe initiative, Yania Ondaro-Martinez and Yan Shi at the Miami Project, and Vance Lemmon for helpful discussion.

---

**Figure 1 - figure supplement 1:**
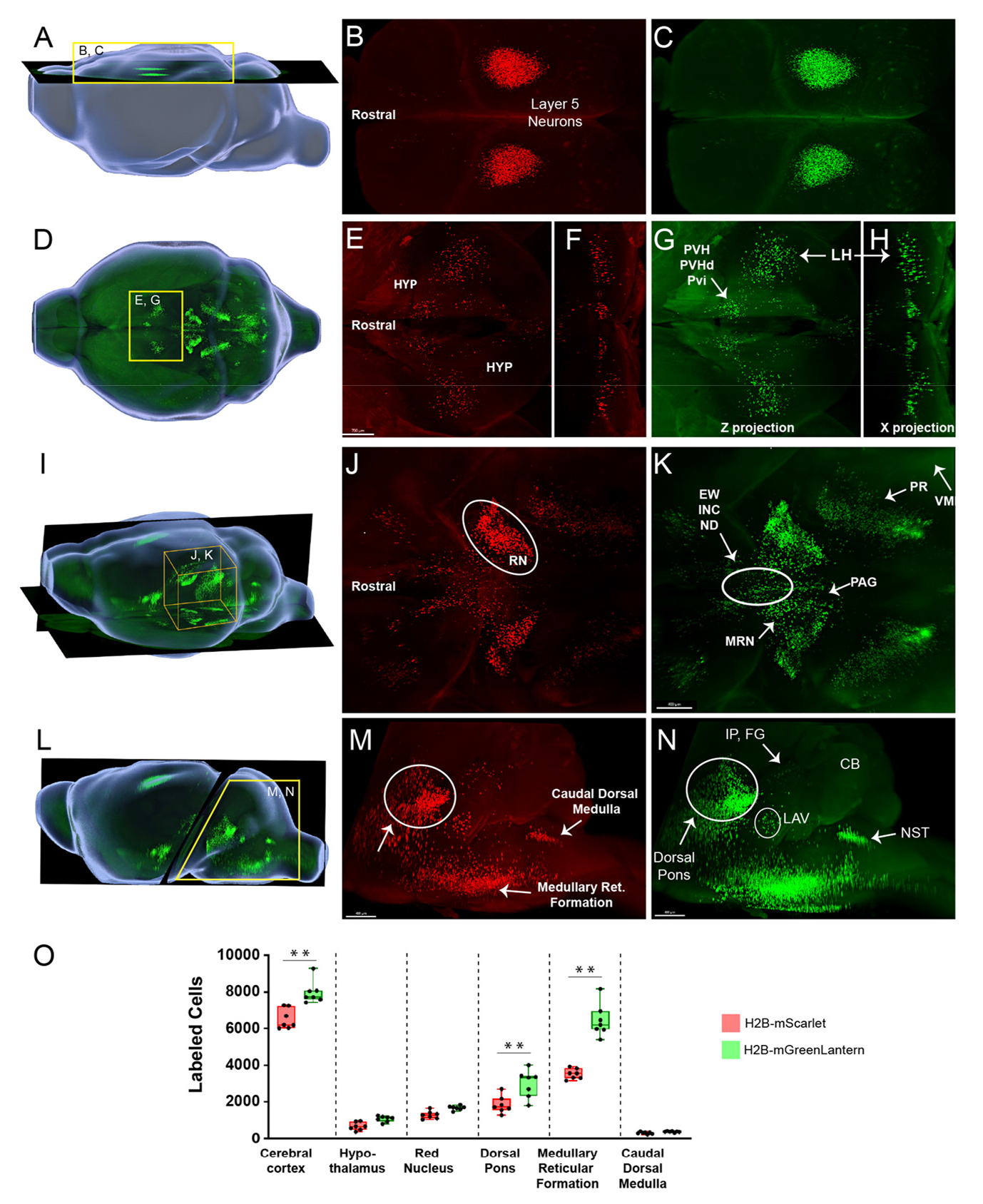
Comparison of supraspinal labeling by nuclear localized mScarlet and mGreenLantern. Whole brains were cleared using 3DISCO and imaged by LSFM two weeks after lumbar co-injection of mixed AAV2-retro-mScarlet and mGreenLantern. **(A)** 3D lateral view of a mouse brain with the plane of image for b and c indicated. **(B,C)** Dorsal view of CST neurons labeled with mSc (B) or mGL (C). (D) Dorsal view of a mouse brain, with hypothalamic nuclei outlined by yellow inset. **(E-H)** Dorsal and coronal views of the hypothalamus with supraspinal neurons labeled by mSc (E,F) or mGL (G,H). (I) Laterodorsal view of the brain with midbrain and rostral pons outlined (yellow box). **(J.K)** Dorsal view of the midbrain and pons with supraspinal neurons labeled by mSc (J) or mGL (K) **(L-N)** Lateral view of the mouse brain with an inset outlining the brainstem and cerebellum and shown as higher magnification images in M for H2BmSc and **N** for H2BmGL **. (O)** Quantification of cells expressing nuclear localized mSc and mGL in six Identified brain regions. Detection of mGL was significantly higher in cortex, pons, and the medullary reticular formation (**p<.01,2-WAY ANOVAwith post-hoc Sidak’s). N=7 animals. Scale bar, **E,G** 700 pm; **J,K** 400 pm; **m,n,** 400 pm. HYP, hypothalamus; PVH, paraventricular hyp. nucleus; PVHd, Paraventricular hyp descending; LH, Lateral hypothalamus; VMH, ventromedial hyp. Nucleus; RN, red nucleus; PR, pons reticular formation; PAG, periqueductal gray; MRN, midbrain reticular nucleus; EW, Edinger-Westphal nucleus; INC, interstitial nucleus of Cajal; ND, nucleus of Darkschewltsch; IP, interposed nucleus; FG, fastigial nucleus; LAV, lateral vestibular nucleus ; NST, nucleus of solitary tract ; SC, spinal cord.

**Figure 1 - figure supplement 2:**
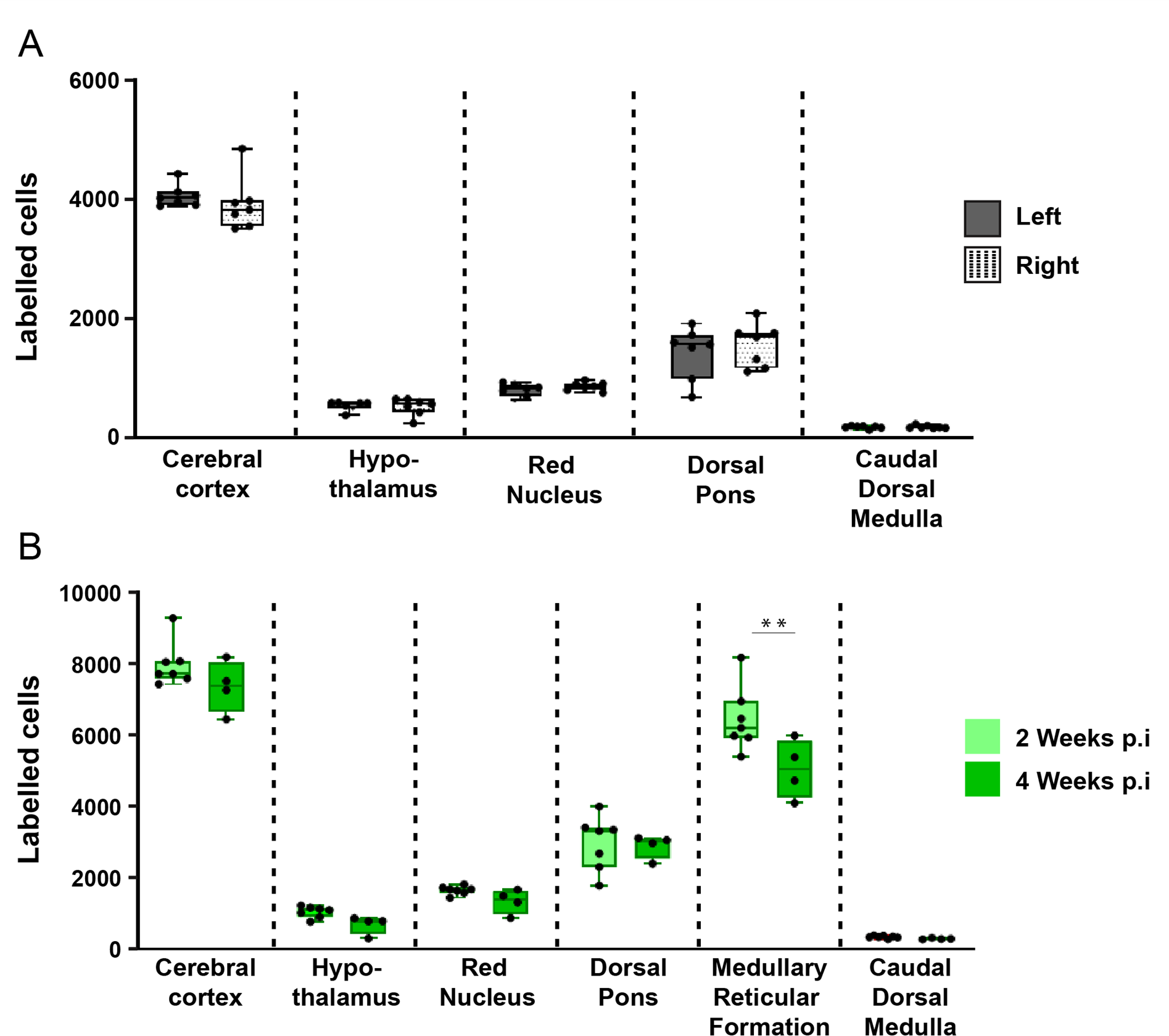
Detection of supraspinal neurons is balanced across the mldllne and maximal by two weeks post-injection. **(A)** Adult mice received lumbar injection of AAV2-Retro-H2B-mGL, followed two weeks later by tissue clearing of brain tissue, LSFM Imaging, and spot detection In lmarls software. The number of detected nuclei did not differ significantly on the right versus left side In any of the analyzed brain regions (p>.05, paired t- test; N= 7 animals). **(B)** Adult mice received lumbar Injection of AAV2-retro-H2B-mGL and survived either two or four weeks, followed by brain tissue clearing, LSFM Imaging, and spot detection In lmarls software. The number of detected nuclei was not significantly elevated In any of the analyzed brain regions at four weeks, compared to two weeks (p>.05, one-tailed t- test; N=7 at 2 weeks, 4 at 4 weeks).

**Figure 2 - figure supplement 1:**
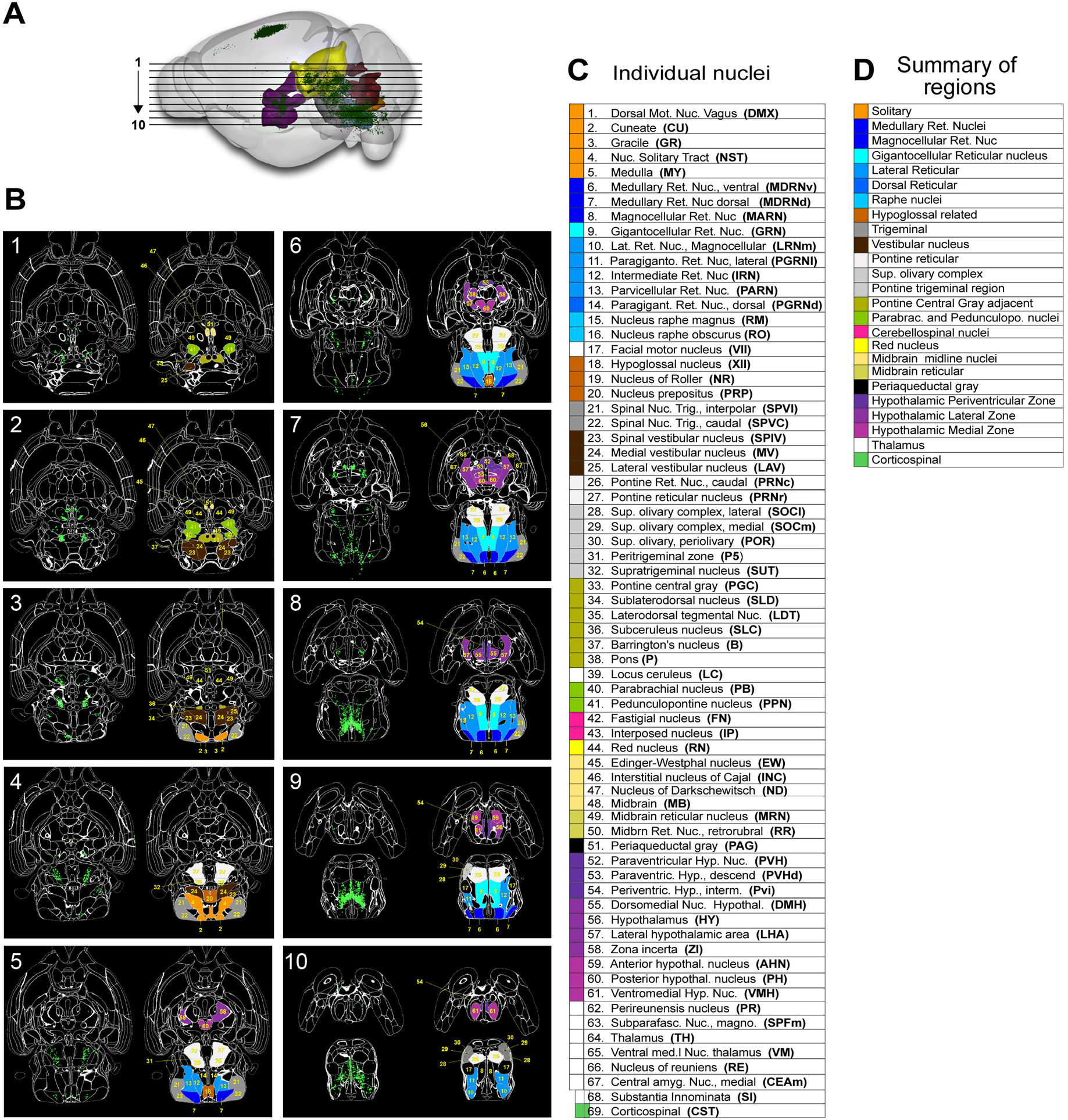
3D imaging follwed by image registraion and nuclei detection by a Map and Cellfinder identifies brain regions containing Supraspaial prijection neurons. Brainrender visualization of a mouse brain after lumbar Injection of AAV2-retro-H2B-mGL, 3D Imaging, registration, and nuclei detection. Detected nuclei are shown In green and selected brain regions Indicated by color. Horizontal lines indicate the planes of images presented in b. **(B)** Horizontal sections produced by Cellfinder, with detected nuclei shown in green, identified brain regions outlined in white, and supraspinal regions coded by colors and numbers corresponding to the key shown in C. Section 1 is most dorsal and section 10 most ventral. **(C)** A key that links color and number to the full name and abbreviation of sixty-nine identified supraspinal brain regions. **(D)** A list of summary regions, in which adjacent supraspinal regions are combined for simplified analysis.

**Figure 4 -figure supplement 1.**
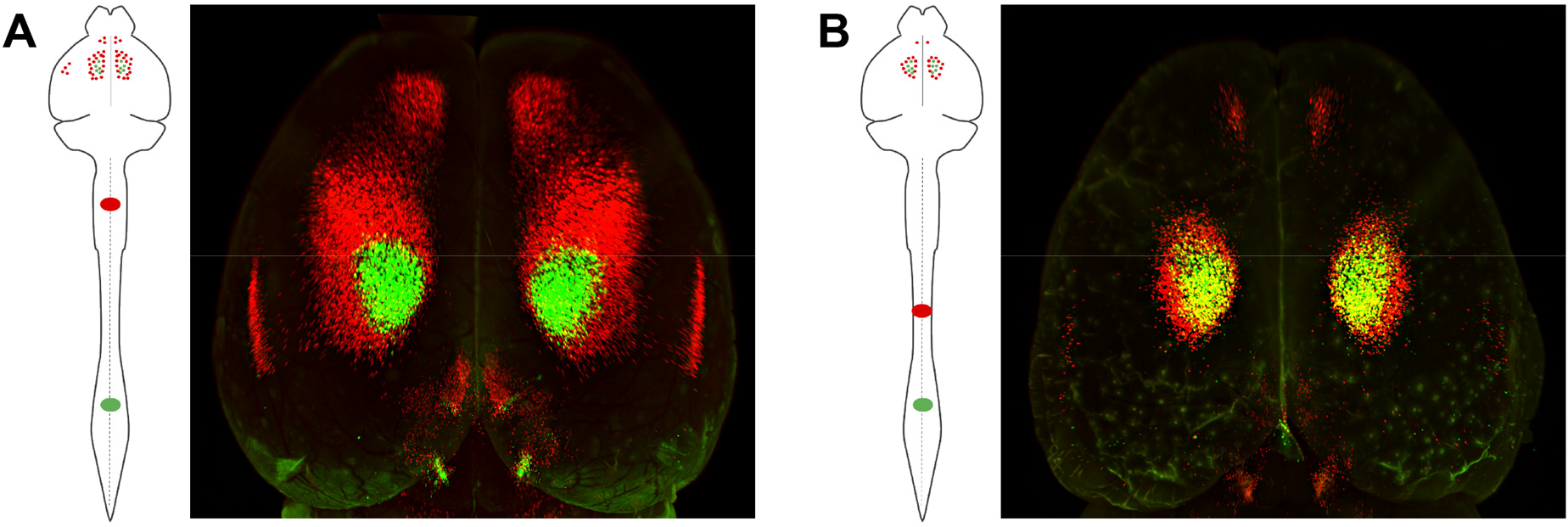
Topographic mapping of the corticospinal projection Is arranged as concentric rings with caudal projections Interior. **(A)** Adult mice received lumbar (L1) Injection of AAV2-retro-mGI and 04 Injection of AAV2-retro-mSc, followed two weeks later by tissue clearing and Imaging. A dorsal view of cortex shows lumbar label (green) completely surrounded by cervical (red). **(B)** Adult mice received lumbar (L1) Injection of AAV2-retro-mGI and thoracic (T10) Injection of AAV2-retro- mSc, followed two weeks later by tissue clearing and Imaging. A dorsal view of cortex shows lumbar label (green) surrounded by a narrow rim of thoracic label. Comparing Ato B Indicates a pattern of concentric zones of projection to progressively more caudal targets In the spinal cord.

**Figure 6 - figure supplement 1.**
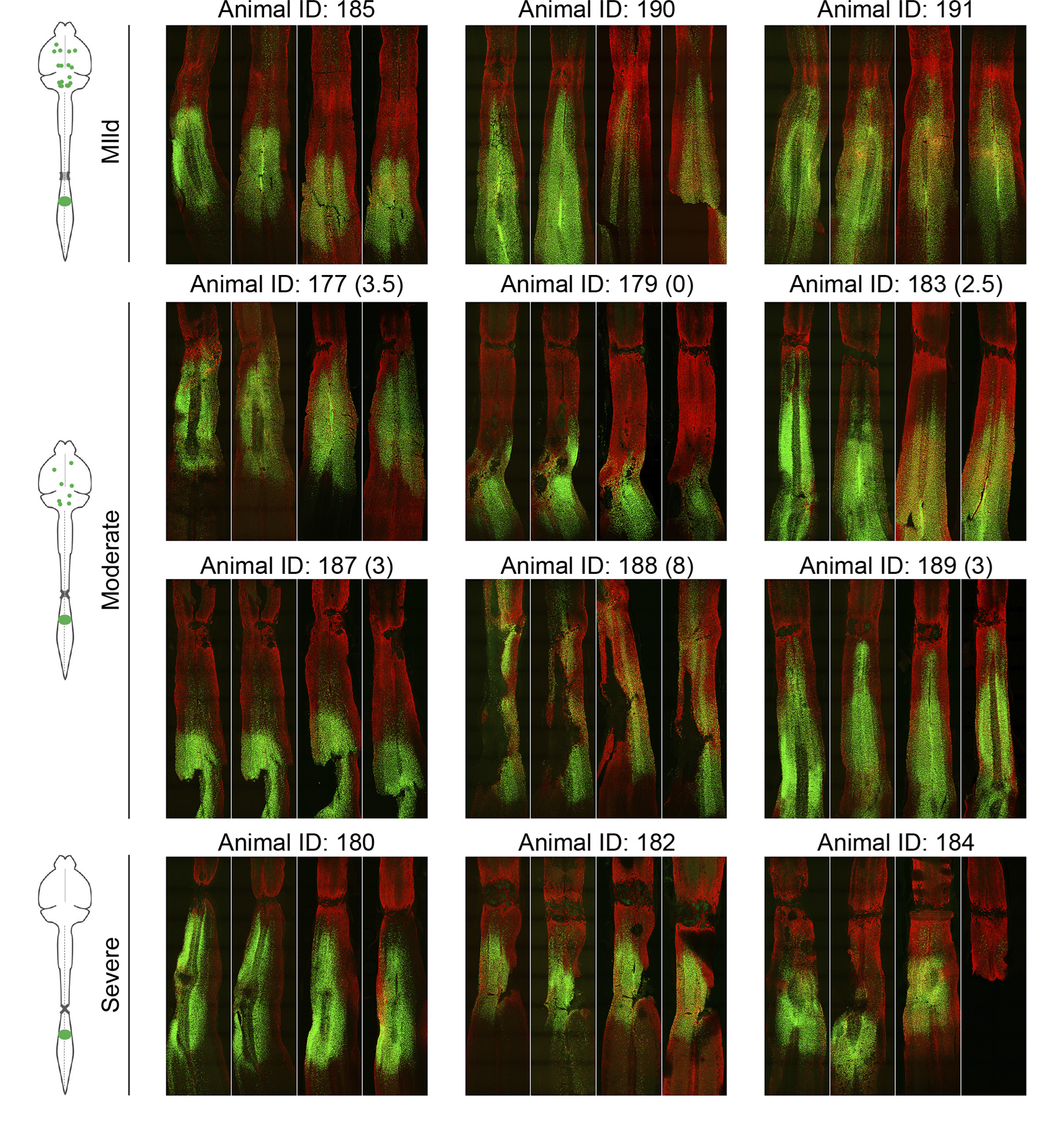
Horizontal spinal sections with GFAP labeling for all injured animals. Adult mice received crush injuries to T11 spinal cord using forceps with stoppers of defined width to control the severity. Seven weeks later AAV2-retro-H2B-mGL was injected to L1 spinal cord, and two weeks post-injection animals were sacrificed and a complete series of 100μm horizontal sections of spinal cord prepared. Immunohistochemistry for the astrocytic marker GFAP (red) defines the site of injury, and H2B-mGL signal (green) indicates neurons that took up the virus. Note that after mild injury the GFAP signal is largely continuous across the crush site, whereas after severe injuries a GFAP-negative region is evident, indicating the absence of astrocytic bridges.

**Figure 7 - figure supplement 1.**
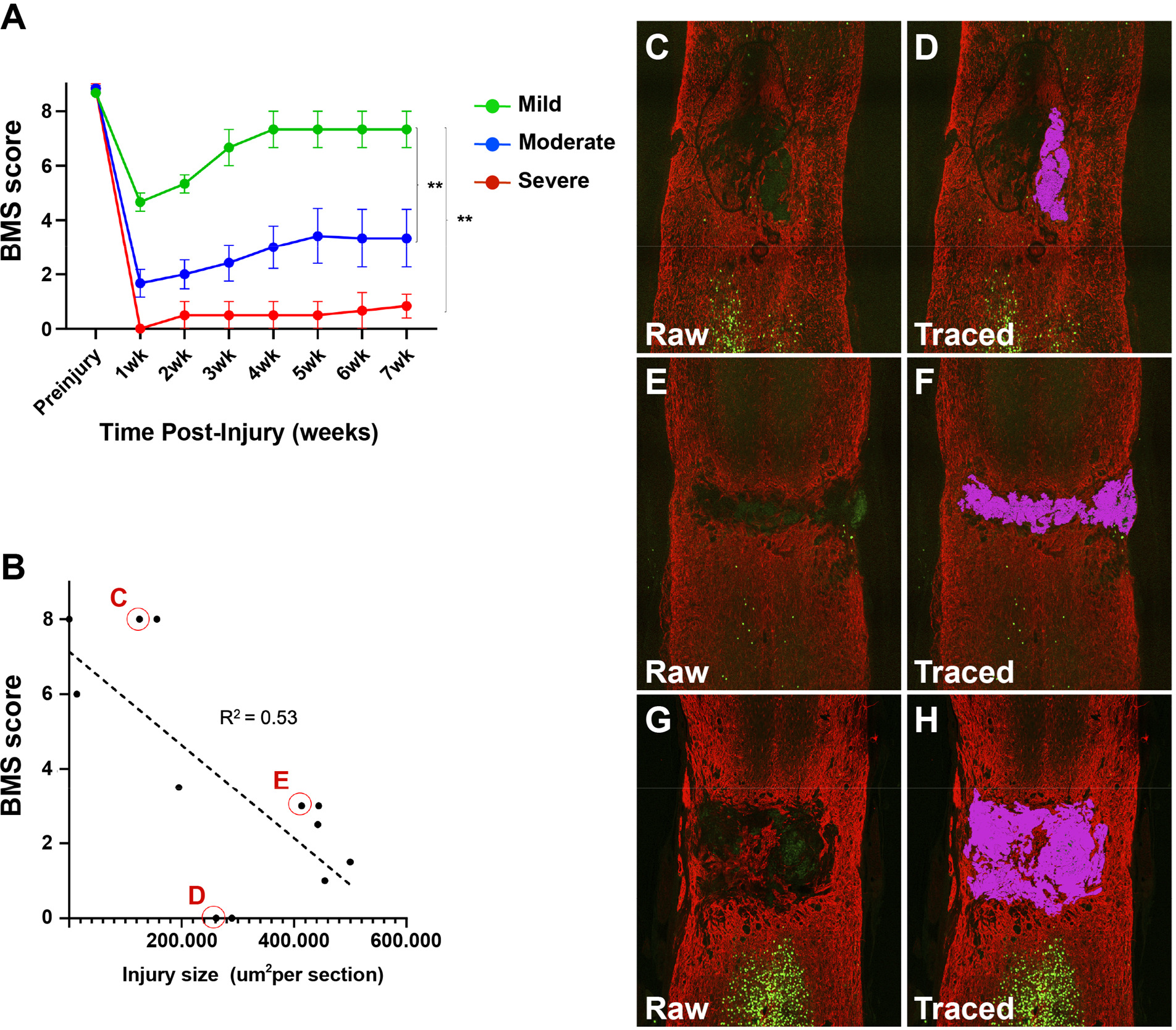
Locomotor recovery from spinal Injury correlates partially with lesion size. **(A)** BMS scores, measured weekly for seven weeks, from mice that received thoracic crush Injury of three different severities. **(B)** Correlation between BMS score at seven weeks and the average injury area measured in spinal cord sections. Circled datapoints correspond to example images to the right. Note that datapoint H has a better motor score than F despite the larger injury size. **(C-H)** Example images of spinal cord stained with GFAP. Injury areas were defined as GFAP-negative cavities, indicated in purple on panels E, F, and H. ** p<.01 Repeated measures ANOVA. N=12 animals: 3 moderate, 6 moderate, and 3 severe

**Figure 7 - figure supplement 2.**
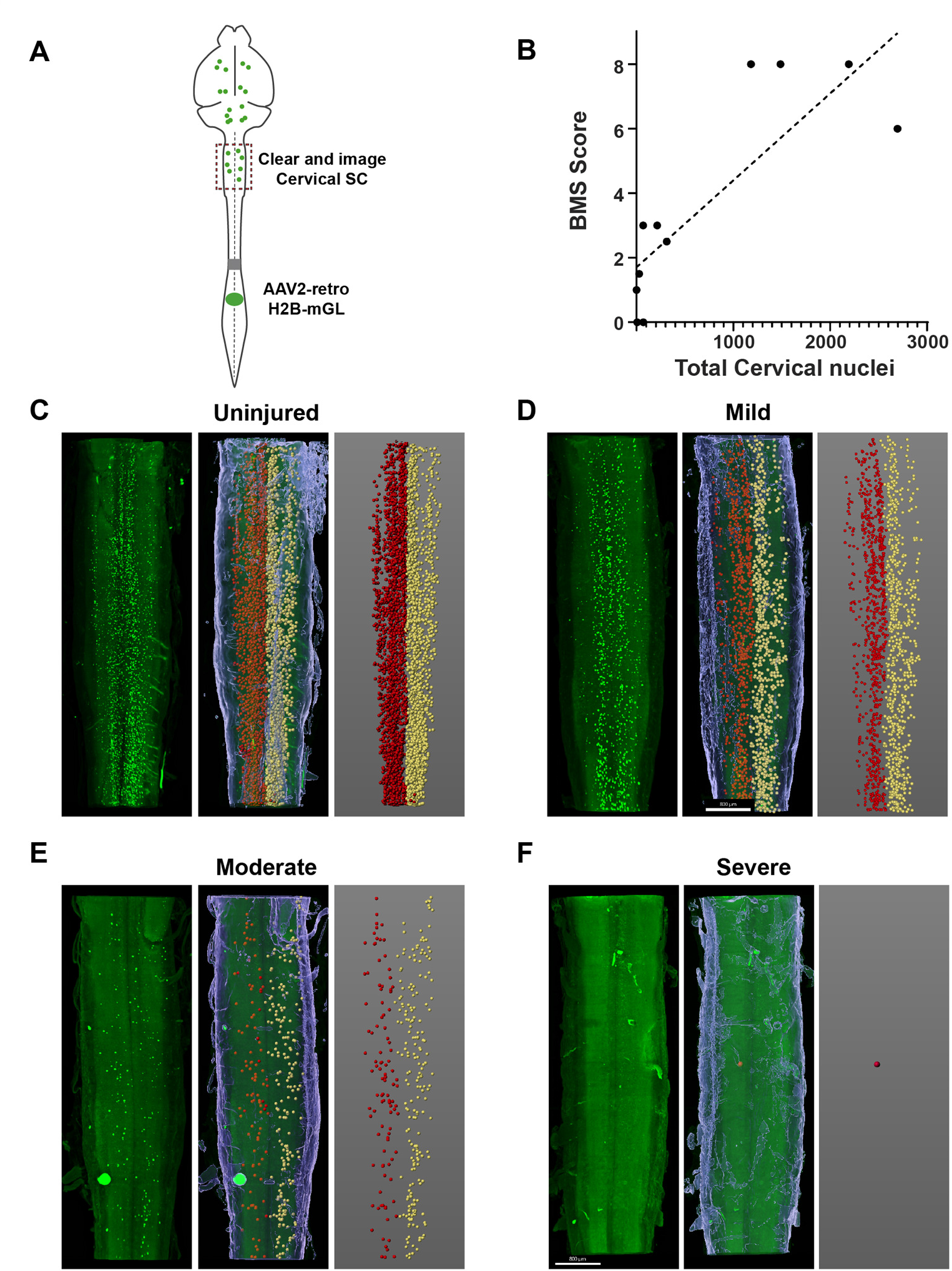
The number of spared propriospinal neurons in cervical spinal cord correlates with locomotor recovery after spinal injury. **(A)** Experimental design. Animals received spinal crush injuries of controlled width, followed by weekly BMS testing for seven weeks, then lumbar injection of AAV2-retro-H2B-mGL. Along with brain clearing and imaging described elsewhere, cervical spinal cord was also cleared and imaged. **(B)** Linear regression between BMS score at seven weeks post-injury and the number of spared neurons in cervical spinal cord (R^2^ = 0.68, slope differs significantly from zero, p<.01 simple linear regression). **(C-F)** Example images of cervical spinal cord. Left panels show raw images, right panels show the detection of labeled nuclei by Imaris with left propriospinal neurons in red and right propriospinal neurons in yellow, and the middle panels show the overlay.

